# PRDM proteins control Wnt/β-catenin activity to regulate craniofacial chondrocyte differentiation

**DOI:** 10.1101/2021.05.21.445211

**Authors:** Lomeli C. Shull, Hyun Min Kim, Ezra S. Lencer, Susumu Goyama, Mineo Kurokawa, James C. Costello, Kenneth Jones, Kristin B. Artinger

## Abstract

Cranial neural crest (NCC)-derived chondrocyte precursors undergo a dynamic differentiation and maturation process to establish a scaffold for subsequent bone formation, alterations in which contribute to congenital birth defects. Here, we demonstrate that transcription factor and histone methyltransferase proteins Prdm3 and Prdm16 control the differentiation switch of cranial NCCs to craniofacial cartilage. Loss of either results in hypoplastic and unorganized chondrocytes due to impaired cellular orientation and polarity. We show that PRDMs regulate cartilage differentiation by controlling the timing of Wnt/β-catenin activity in strikingly different ways: *prdm3* represses while *prdm16* activates global gene expression, though both by regulating Wnt enhanceosome activity and chromatin accessibility. Finally, we show that manipulating Wnt/β-catenin signaling pharmacologically or generating *prdm3-/-;prdm16-/-* double mutants rescues craniofacial cartilage defects. Our findings reveal upstream regulatory roles for Prdm3 and Prdm16 in cranial NCCs to control Wnt/β-catenin transcriptional activity during chondrocyte differentiation to ensure proper development of the craniofacial skeleton.

**Highlights:**

1. Prdm3 and Prdm16 are required for chondrocyte organization in vertebrate craniofacial cartilage
2. Loss of Prdm3 and Prdm16 alters expression of Wnt/β-catenin signaling components
3. Prdm3 and Prdm16 oppositely control global chromatin accessibility
4. Prdm3 and Prdm16 cartilage defects can be rescued pharmacologically or genetically

## Introduction

The formation of the craniofacial skeleton relies on precise cellular and molecular processes across multiple tissues and cell types to facilitate cranial neural crest (NCC) development [1-3]. The cranial NCCs that give rise to chondrocytes will form cartilaginous structures that will lay the foundation for the craniofacial skeleton. Fundamental to the growth of the cartilage is the organization and positional orientation of chondrocytes during proliferation and growth which facilitates proper transitions toward hypertrophy, and the subsequent recruitment of osteoblasts to deposit bony matrices over or around the cartilaginous template [4-8]. These processes require extensive temporal and spatial regulation as proper formation of these cartilage nodules establishes the cartilage template that will be used as a scaffold for subsequent bone formation. Alterations to the gene regulatory networks (GRNs) and signaling modules that control craniofacial chondrocyte maturation can greatly impact the development of these cartilage elements. Changes to the development of these cartilage scaffolds can negatively affect the formation and function of subsequent bone structures and contribute to craniofacial defects including, but not limited to mandibular hypoplasia, cleft lip/palate, and middle ear defects[9-18].

Alongside GRNs that facilitate chondrogenesis, several key signaling pathways are important for mediating and maintaining chondrocyte development. Canonical Wnt/β-catenin signaling plays a highly dynamic role in facilitating chondrocyte differentiation. Early pre-chondrocyte cell-cell contacts and intercalations are regulated by N-cadherin, α-and β-catenin within pre-cartilage condensations [19-25]. These interactions at cell-cell junctions are maintained as more stable cartilage structures are established, but then gradually destabilized as chondrocytes mature [8, 19-22, 26-29]. Alongside maintenance of cell-cell contacts, transcriptionally active nuclear β-catenin also plays a role in facilitating chondrocyte maturation. While required early in prechondrogenic cells, β-catenin is gradually downregulated as chondrocytes become more differentiated [19, 23-25, 30-32]. β-catenin is active again at late stages of chondrocyte hypertrophy to promote osteoblast differentiation and mineralization. However, abnormally high levels of β-catenin at later stages of chondrogenesis preceding hypertrophy are inhibitory to differentiation leading to abnormal hypertrophy, disorganization of chondrocytes and instead stimulate osteoblast differentiation and premature mineralization [23-25, 30-32]. Thus, gradients of Wnt/β-catenin must be balanced to facilitate proper chondrocyte differentiation and maturation.

Crucial to the transcriptional activity of β-catenin is nuclear translocation of β-catenin and assembly of the Wnt/β-catenin enhanceosome transcriptional complex consisting of Bcl9/Bcl9l, Pygo1/2, Lef1/Tcf, and chromatin modifiers such as Brg1 (Smarca4a), CBP/p300, among others [33, 34]. Of these, Bcl9 plays a versatile role in facilitating β-catenin cellular localization and nuclear transcriptional activity, as Bcl9 can shuttle between the cytoplasm and nucleus. Bcl9 can compete with α-catenin, precluding β-catenin’s interaction with α-catenin at adhesions complexes [34-39]. Once nuclear, Bcl9 acts as the main scaffolding protein necessary for nuclear retention of β-catenin along with Bcl9’s binding partner, Pygo1/2 [38, 39]. Together Bcl9 and Pygo1/2 assemble other chromatin remodelers and transcription factors that then drive Wnt/β-catenin transcriptional activity [38, 40-42]. While the processes governing cranial NCC-derived chondrocyte differentiation during craniofacial development have well been characterized, the upstream mechanisms driving spatial and temporal activation and repression of GRNs and signaling pathways, in particular canonical Wnt/β-catenin and the formation of the Wnt enhancesome complex, remain largely unknown.

The role of chromatin modifiers in cranial neural crest development and the formation of the craniofacial skeleton have only recently become of interest due to the identification of several chromatin remodelers and their association with craniofacial disorders [43]. Among these regulatory factors that are important for cranial neural crest development and formation of the craniofacial skeleton are the PRDM (Positive Regulatory Domain) family of lysine methyltransferases. These modulators control gene expression through several different mechanisms: by epigenetically modulating chromatin at specific target gene promoters, by directly binding DNA via zinc finger domains, or by interacting with other protein complexes and histone modifying enzymes to control gene expression [44-46]. Human genome wide association studies have associated SNPs in the genes encoding two of these PRDMs, PRDM3 (EVI1/MECOM) and PRDM16, to craniofacial abnormalities including cleft palate and influencing variation in facial morphology [47-51].

Traditionally, while PRDM3 and PRDM16 have been associated with transcriptional repression by their ability to enzymatically modify histone H3 lysine 9 (H3K9), a mark associated with transcriptional repression, more recently been involved in gene activation through methylation of histone 3 lysine 4 (H3K4) [52-54]. PRDM3 and PRDM16 can indirectly regulate gene expression by forming complexes with other co-factors and chromatin modifying enzymes [45, 55-59]. Putative DNA binding motifs for PRDM3 and PRDM16 have further implicated their potential to also directly bind DNA to mediate gene expression [44-46]. PRDM3 and PRDM16 are important in a variety of developmental processes in mouse and zebrafish, acting as both repressors and activators of specific GRNs depending on cellular context [54, 60-71]. We and others have previously shown loss of *Prdm3* and *Prdm16* in zebrafish and mice causes craniofacial defects including hypoplasia of cartilage elements [61, 72]. Importantly, while a level of genetic compensation exists between these two paralogs as a result of their high amino acid sequence homology and similar developmental expression patterns during embryogenesis, there is evidence suggesting they also have diverging roles from each other [72]. Recent studies have implicated the importance of *Prdm3* and *Prdm16* in craniofacial development, however, the exact mechanism(s) of their action in mediating proper cranial NCC-derived cartilage differentiation and the proper formation of the craniofacial skeleton remain largely unknown.

In this study, we utilized zebrafish and mouse genetic models to dissect the molecular functions of Prdm3 and Prdm16 in chondrogenesis during craniofacial development. Based on our data, we hypothesize under normal developmental conditions, Prdm3 acts as a transcriptional repressor while Prdm16 serves as an activator of similar gene expression targets in cranial NCCs to regulate chondrocyte differentiation by controlling canonical Wnt/β-catenin transcriptional activity during chondrocyte differentiation to ensure proper spatial and temporal development of the zebrafish and mouse craniofacial skeleton.

## Results

### *prdm3* and *prdm16* are required for craniofacial chondrocyte stacking and polarity

Previous studies have shown that despite genetic compensation between *prdm3* and *prdm16*, and a third PRDM family member, *prdm1a*, the individual loss of *prdm3* or *prdm16* causes moderate overall craniofacial phenotypes in the developing zebrafish pharyngeal skeleton [72] (**Fig. 1A-1C, 1A’-1C’**). However, significant hypoplasia of cartilage elements within the craniofacial skeleton suggested changes to the subcellular development of these structures. To visualize cartilage and bone, *prdm3-/-* and *prdm16-/-* larvae were stained with Alcian blue and Alizarin red. High magnification of the cartilage elements within all areas of the skeleton revealed changes in chondrocyte-to-chondrocyte cellular adhesions and contacts (**Fig. 1D**). Unlike the organized, stacked chondrocytes of wildtype embryos, chondrocytes in *prdm3-/-* and *prdm16-/-* mutants were highly disorganized. This was quantified by measuring the angle between three adjacent chondrocytes in the direction of growth in the ceratobranchial cartilage elements (**Fig. 1E-1F**). The more organized the cells, the closer the angle between adjacent cells reaches 180°. Loss of *prdm3* and *prdm16* caused a significant (33.4% and 34.6%, respectively) reduction of the angle between adjacent cells (**Fig. 1F**). The number of cells per unit of area within these cartilage structures at 6 dpf was increased in both *prdm3-/-* (∼32.3% increase) and *prdm16-/-* (∼29.6% increase) mutants (**Fig. 1G**). Previous work has demonstrated that outgrowth of pre-cartilaginous condensations requires cell-cell intercalations and changes in cell shape (hypertrophy) rather than through extensive proliferation [4, 7, 8]. Loss of *prdm3* and *prdm16* causes no change in proliferation during early development of cartilage structures [72]. As such, the increased number of cells per tissue area observed in *prdm3* and *prdm16* mutants likely results from the failure of these chondrocytes to intercalate properly and their inability to expand their cell shape for growth. To further evaluate changes in cell intercalation and extension, live imaging from 56 hpf to 72 hpf was performed on wildtype, *prdm3-/-* and *prdm16-/-* crossed into the Tg(*sox10:EGFP*) transgenic background. Live imaging revealed failure of the chondrocytes within cartilage structures to extend during development in both *prdm3* and *prdm16* mutants (**Supplemental movies 1-3 and Supplemental Fig. 1A-1C**). In addition to the cellular changes observed morphologically, these results also suggested chondrocyte cell polarity was also abrogated in *prdm3* and *prdm16* mutants.

**Figure 1.**
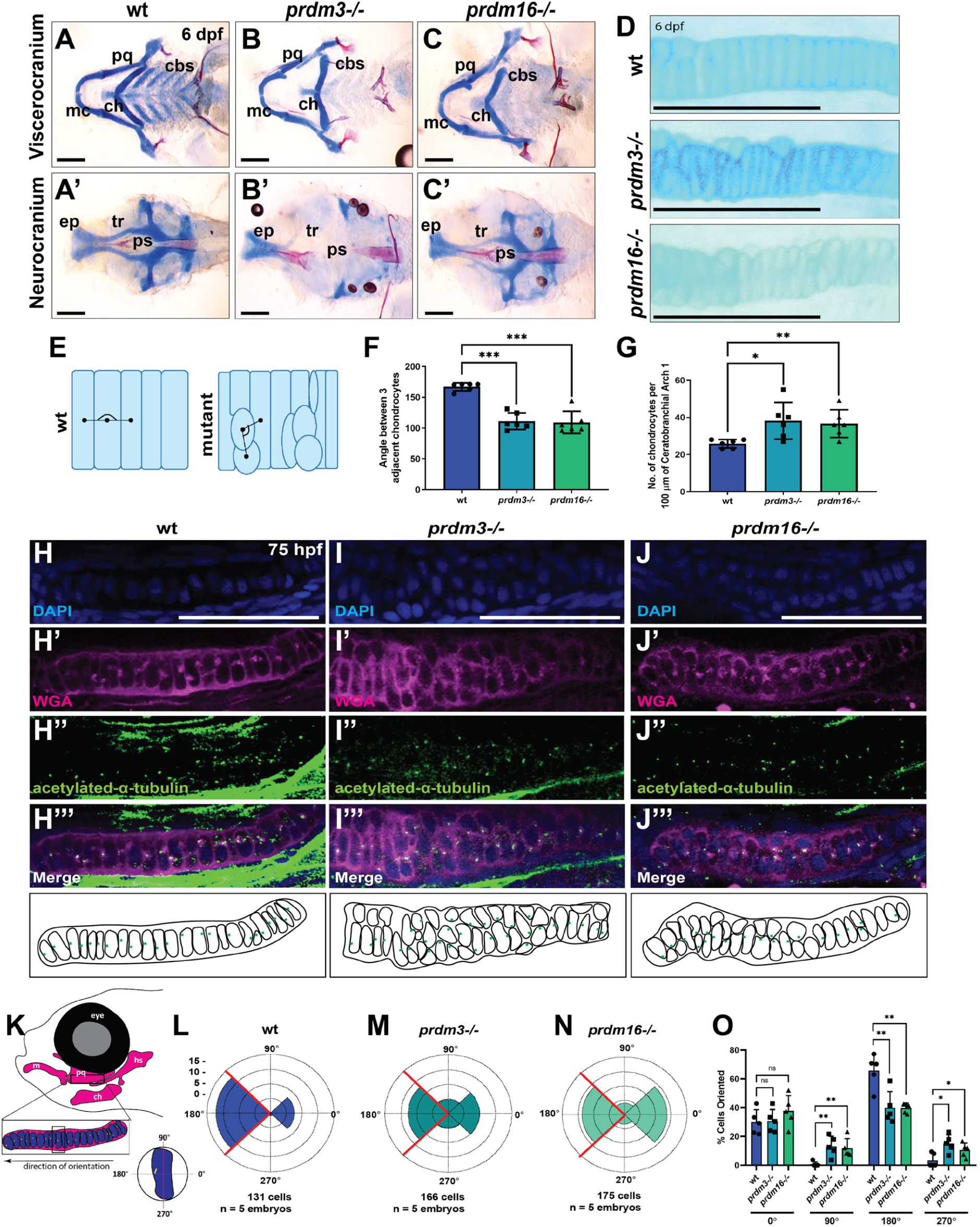
*prdm3* and *prdm16* are necessary for chondrocyte stacking and polarity in the zebrafish craniofacial skeleton. (A-G) Wildtype (wt), *prdm3-/-, prdm16-/-* zebrafish embryos were collected at 6 days post fertilization (dpf) and stained for Alcian blue and Alizarin red. Images of dissected and flat mounted viscerocranium (A-C) and neurocranium (A’-C’). (D) High magnification of chondrocytes in the cartilage elements shows altered chondrocyte stacking with loss of *prdm3* and *prdm16*. (E-F) Quantification of chondrocyte stacking and organization as schematized in (E). Loss of *prdm3* and *prdm16* significantly reduces the angle between adjacent chondrocytes compared to wildtypes (F) (n = 6 individuals for each genotype). (G) Quantification of the number of chondrocytes per 100 µm of a cartilage element shows an increase in cell number in *prdm3* and *prdm16* mutants (n = 6 individuals per genotype). cbs, ceratobranchials; ch, ceratohyal; ep, ethmoid plate; mc, Meckel’s cartilage; pg, palatoquadrate; ps, parasphenoid; tr, trabeculae. Scale bar, 100 µm. (H-O) Wildtype, *prdm3-/-* or *prdm16-/-* larvae were immunostained with acetylated alpha-tubulin to label microtubule organizing centers (MTOCs) at 75 hpf (H”-J”). Larvae were counter stained with DAPI (H-J) and Wheat Germ Agglutinin (WGA) (H’-J’). Shown are lateral high magnification views of the developing palatoquadrates. Scale bar, 50 µm. (K-O) Quantification of chondrocyte polarity by noting the positioning of acetylated alpha-tubulin puncta, as depicted in (K). Positioning was tracked through z-stack images and the number of acetylated alpha-tubulin puncta in each quadrant for each cell assessed were tabulated for wildtype (L), *prdm3-/-* (M), and *prdm16-/-* (N). (O) Percentage of cells at each indicated quadrant normalized to the total number of cells for each genotype. (n = 5 for each genotype). Scale bar, 100 µm. * p ≤ 0.05, ** p ≤ 0.005, *** p ≤ 0.001, ns, not significant, Student’s *t* test.

To further assess changes in cell polarity, wildtype, *prdm3-/-* or *prdm16-/-* larvae were stained with acetylated-alpha tubulin to label microtubule-organizing centers (MTOCs) at primary cilia at 75 hpf, a time point after which chondrocyte cell-cell rearrangements and cartilage polarity have stabilized and cells are oriented in a specific direction for growth. During cartilage morphogenesis, positions of MTOCs undergo dynamic rearrangement during periods of cell-cell intercalation (48-54 hpf) [8]. Initially, MTOCs in pre-chondrocytes are oriented towards the center of each condensation of the cartilage. As developmental time progresses, these cells rearrange to orient themselves in a uniform manner along dorsal-ventral axes and by 66 hpf, MTOC positions have stabilized [8]. At 75 hpf, the MTOCs in the chondrocytes of the palatoquadrate of wildtype larvae were localized ventrally toward the Meckel’s cartilage and the jaw joint junction (**Fig. 1H-H’’’**). However, the MTOCs in stage-matched *prdm3-/-* (**Fig. 1I-1I’’’)** and *prdm16-/-* (**Fig. 1J-1J’’’)** failed to rearrange and orient themselves uniformly along this ventral axis and instead were positioned dorsally, or toward the center of the original condensation (**Fig. 1I-1I’’’, 1J-1J’’’**). Quantification of MTOC orientation in the palatoquadrate showed 65% of chondrocytes at this stage were ventrally polarized, as denoted by MTOC puncta positioned directionally at the 180° quadrant of the cell (**Fig. 1K-1L, 1O**). Conversely, loss of *prdm3* and *prdm16* caused a significant reduction (∼25%) in the percentage of uniformly ventrally oriented chondrocytes (180°), which corresponded to an increase in the number of chondrocytes that were instead positioned dorsally (0°) or toward the center of the cartilage element (90° or 270°) (**Fig. 1M-1O**). Taken together, these results demonstrate *Prdm3* and *Prdm16* are important for facilitating proper chondrocyte differentiation including orientation, intercalation and organization in the cartilage elements that form during craniofacial cartilage morphogenesis.

### Neural crest specific function of *Prdm3* and *Prdm16* is required for chondrocyte organization in Meckel’s cartilage of the developing murine mandible

We have recently shown that while Prdm3 and Prdm16 have divergent roles across mammalian and non-mammalian vertebrates, there is also evidence of conserved functions across vertebrate species [72]. Prdm3 and Prdm16 have high protein abundance in the facial prominences during mouse craniofacial development (**Supplemental Fig. 2A-2B, 2A’-2B’**).To evaluate changes in chondrocyte development during the formation of the mammalian craniofacial skeleton with loss of *Prdm3* and *Prdm16*, we conditionally ablated *Prdm3* and *Prdm16* in the neural crest lineage using the *Wnt1-Cre* driver [10, 73]. Both homozygous *Prdm3*^*fl/fl*^*;Wnt1-Cre*^*+/Tg*^ and *Prdm16*^*fl/fl*^*;Wnt1-Cre*^*+/Tg*^ mutant animals were born at Mendelian ratios and survived to E18.5. We did not assess the viability of mutant embryos past E18.5 but predict, based on previous studies, that both mutants would not survive postnatally due to either defects in heart development (*Prdm3*) or failure to thrive due to cleft palate defects (*Prdm16*) [60, 70,72, 74, 75]. Homozygous *Prdm3* mutant animals developed an overall subtle craniofacial phenotype, namely mild anterior mandibular hypoplasia and slight defects in snout extension (**Fig. 2B-2B”, 2E**). Homozygous *Prdm16*^*fl/fl*^*;Wnt1-Cre*^*+/Tg*^ mutants present with a variety of craniofacial defects, similar to those observed with loss of *Prdm16* in the early epiblast suggesting a cell autonomous function in the neural crest [60, 72]. Defects observed include snout extension defects and anterior mandibular hypoplasia, as well as a secondary cleft palate and middle ear defects with severe hypoplasia of the tympanic rings, incus and malleus and retroarticular process of the squamosal bone (**Fig. 2C-2C’’, 2E, Supplemental Fig. 2C-2E, 2F-2H, 2F’-2H’, 2F’’-2H’’**).

**Figure 2.**
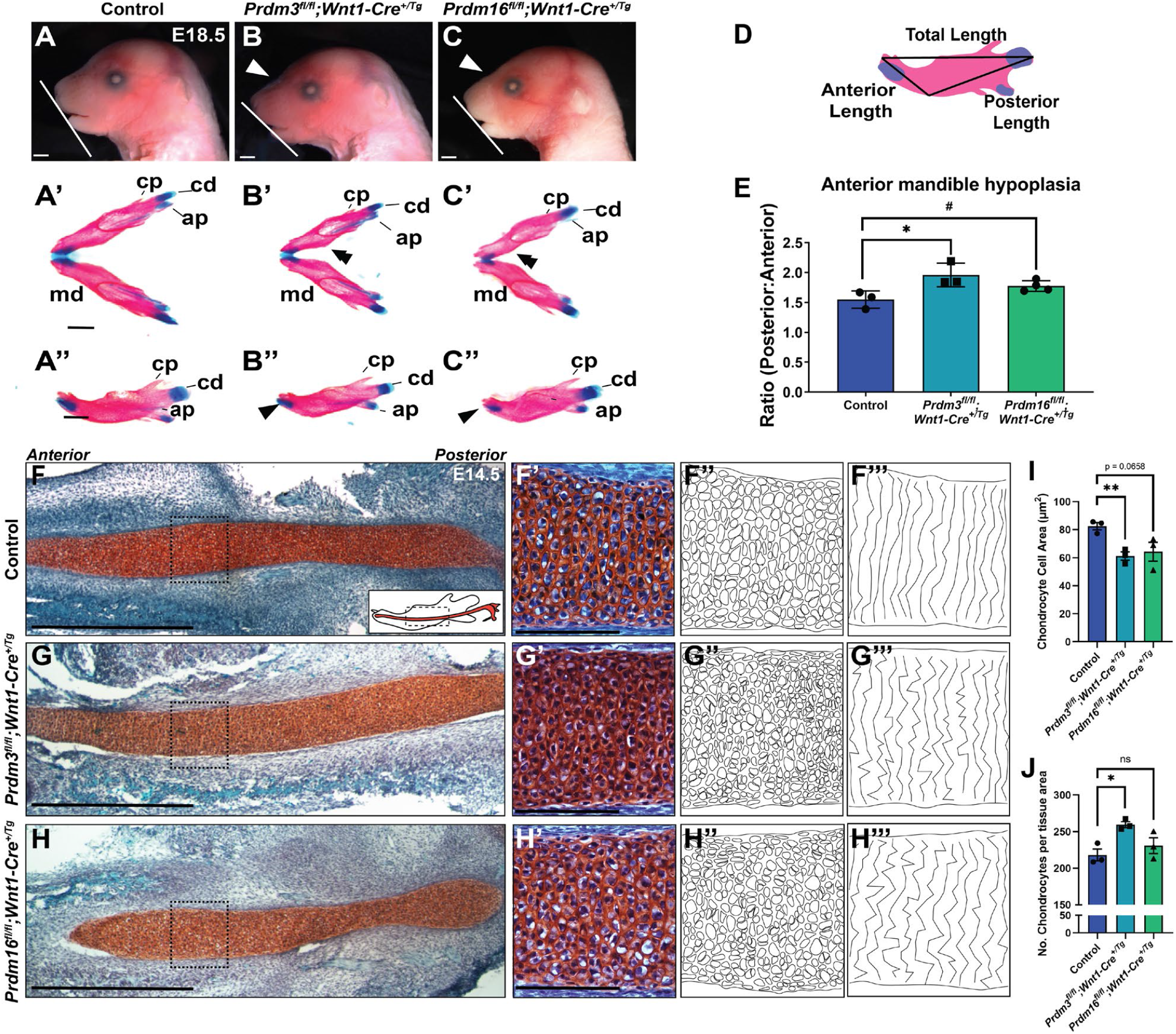
Loss of *Prdm3* and *Prdm16* in the murine neural crest lineage alters chondrocyte organization in the Meckel’s cartilage of the developing mandible. (A-M) *Prdm3*^*fl/fl*^ or *Prdm16*^*fl/fl*^ females were bred for timed matings to *Prdm3*^*fl/+*^*;Wnt1-Cre*^*+/Tg*^ or *Prdm16*^*fl/+*^*;Wnt1-Cre*^*+/Tg*^ males and embryos were collected at the indicated timepoints. (A-C) Lateral images of head show snout defects (white arrowhead) and hypoplasia of the mandible (white line) in both *Prdm3* and *Prdm16* mutants at E18.5. (A’-C’) Alcian blue and Alizarin red stained mandibles were dissected and removed from control or mutant animals. (A’’-C’’) Lateral views of the right half of the mandible. (D-E) Quantification of anterior mandibular hypoplasia was performed by measuring the ratio between the length of the anterior and posterior portions of the mandible (P:A) as schematized in (D). (F-H) Safranin O and Fast Green stained sagittal sections of the Meckel’s cartilage at E14.5 in control (F), *Prdm3* (G) or *Prdm16* (H) mutants. Scale bar, 100 µm. (F’-H’) High magnification of the chondrocytes from the boxed regions in (F-H), Scale bar, 100µm. (F’’-H’’) Schematized cellular arrangements of chondrocyte cell shapes. (F’’’-H’’’) Schematic illustrating chondrocyte organization; lines were drawn connecting nuclei from the top to the bottom of the Meckel’s cartilage section. (I-J) Quantitation of chondrocyte cell area (I) and cell number per tissue area (J) on n = 3 for each genotype. ap, angular process; cd, condylar process; cp, coronoid process; md, mandible. * p ≤ 0.05, Student’s *t* test.

The consistent and shared phenotype across both *Prdm3* and *Prdm16* mutants was mandibular hypoplasia (**Fig. 2A-2E**). Alcian blue and Alizarin Red stained skeletal preparations confirmed the observed mandibular hypoplasia in both *Prdm3* and *Prdm16* mutants (**Fig. 2A’-2C’, 2A’’-2C’’**). The anterior growth of the mandible was significantly reduced in mutants, as quantified by measuring the ratio of the anterior portion of the mandible relative to the posterior length (**Fig. 2D-2E**). This phenotype observed in both *Prdm3* and *Prdm16* mutants parallels some of the hypoplasia seen in *prdm3* and *prdm16* zebrafish mutants and suggests changes to the cellular development and maturation of the chondrocytes in cartilaginous structures (Meckel’s cartilage) that will support the formation of the mandible. To assess the cellular arrangements of the chondrocytes within the developing Meckel’s cartilage histologically, sagittal sections through the Meckel’s cartilages were collected from control, *Prdm3*^*fl/fl*^*;Wnt1-Cre*^*+/Tg*^, and *Prdm16*^*fl/fl*^*;Wnt1-Cre*^*+/Tg*^ at E14.5, the stage of cartilage development where the chondrocytes have undergone rapid growth and are starting to prehypertrophy (**Fig. 2F-2H, 2F’-2H’, 2F’’-2H’’**). Safranin O and Fast Green staining of sections revealed changes to the chondrocyte organization in mutant animals compared to controls. In control animals, chondrocytes were starting to swell and become pre-hypertrophic as evidenced by expansion of the intracellular space (**Fig. 2F’, 2F’’**). These cells were also organized into stacks extending from the top of the cartilage structure to the bottom of the Meckel’s cartilage (**2F’, 2F’’’**). By contrast, *Prdm3* mutant chondrocytes were tightly packed and condensed, suggesting a stall in their maturation process (**Fig. 2G’, 2G’’**). *Prdm16* mutant chondrocytes were unsynchronized, some appeared compacted like those in the *Prdm3* mutants while others seemed to be undergoing accelerated pre-hypertrophy (**Fig. 2H’, 2H’’**). In both *Prdm3* and *Prdm16* mutants, the chondrocytes were not organized in stacks from the top to the bottom of the Meckel’s cartilage (**Fig. 2G’’-2H’’, 2G’’’-2H’’’**). Quantification of the number of cells and the area of chondrocytes within a specified tissue area further supported these cellular changes with a near significant ∼22% decrease in chondrocyte cell area (**Fig. 2I**) and a corresponding ∼16% increase of total number of cells per tissue area (**Fig. 2J**) with loss of *Prdm3* compared to control animals. Decreased cell area was near significance (p = 0.0658) and there were no significant changes in the total number of cells present with loss of *Prdm16*, likely due to heterogeneric differentiation of chondrocytes observed in these animals (**Fig. 2I-2J**). Together, these cellular changes observed in the developing Meckel’s cartilage chondrocytes with loss of *Prdm3* and *Prdm16* suggest altered chondrocyte differentiation which could contribute to the anterior mandibular hypoplasia observed in these mutants. Importantly, the chondrocytes within both zebrafish and mouse craniofacial structures are abnormal with loss of *Prdm3* and *Prdm16* suggesting a conserved function of these factors in regulating chondrocyte differentiation and maturation in vertebrate craniofacial cartilage development.

### *prdm3* and *prdm16* regulate global gene expression in zebrafish cranial neural crest cells

To dissect the molecular mechanisms of *prdm3* and *prdm16* in controlling cranial NCC cartilage derivative differentiation and morphogenesis, we focused on the zebrafish cranial NCCs. Each mutant line was crossed into the *Tg(−4*.*9sox10:EGFP)* background [76]. *sox10:EGFP* positive cranial NCCs were isolated from wildtype, *prdm3-/-* and *prdm16-/-* embryos at 48 hpf via fluorescent activated cell sorting (FACS), harvested for RNA and subjected to RNA-sequencing. Transcriptomic analysis revealed striking differences in overall gene expression with loss of *prdm3* or *prdm16* (**Fig. 3**). In *prdm3-/-*, an overwhelming 2688 genes were significantly upregulated, while only 189 genes were significantly downregulated (**Fig. 3A-3B**). Conversely, in *prdm16* mutants the majority of differentially expressed genes were significantly downregulated (1370), while only 279 were significantly upregulated (**Fig. 3A,3C**). GO Pathway analysis was performed separately on the genes that were upregulated in *prdm3* mutants and those downregulated in *prdm16* mutants. Both analysese revealed enrichment of genes associated with similar biological processes including regulation of histone modifications and chromatin organization, as expected with loss of chromatin modifiers (**Supplemental Fig. 3A-3B)**. Canonical Wnt/β-catenin was identified as the top signaling pathway enriched in both the upregulated genes in *prdm3* mutants and the downregulated genes in *prdm16* mutants (**Fig. 3B-3E, Supplemental Fig. 3C-3D)**. In addition, pathway analysis also associated these differentially expressed genes with maintenance of cell polarity, and cell-cell signaling and cellular components such as apical junction complexes, microtubules and cell-cell adherens junctions (**Supplemental Fig. 3A-3B**). Genes belonging to the canonical Wnt/β-catenin signaling axis including *ctnnb1* and other signal transducers (*fzds, dvls, apc*) were oppositely differentially expressed in *prdm3-/-* and *prdm16-/-* mutants (**Fig. 3B, 3D, Supplemental Fig. 3C-3D**). Downstream Wnt/β-catenin transcriptional targets (*fosab, jun, tcf7, lef1*) were also differentially expressed in both mutants (**Fig. 3B, 3D, Supplemental Fig. 3C-3D**). Interestingly, many of these genes encoded factors involved in the Wnt/β-catenin enhanceosome transcriptional complex (*bcl9/bcl9l, pygo1/2, smarca4a (brg1), ep300a/b, arid1aa/ab, tcf7/lef1*) (**Fig. 3B, 3D, Supplemental Fig. 3C-3D**). These opposing transcriptomic profiles in *prdm3-/-* and *prdm16-/-* mutants were further validated by real-time quantitative polymerase chain reaction (RT-qPCR) on RNA from whole heads of wildtype, *prdm3-/-*, or *prdm16-/-* embryos at 48 hpf (**Fig. 3F-3H**). The trends of gene expression of canonical Wnt effector genes (*ctnnb1, fzd3b, dvl3, apc*) (**Fig. 3F**), components of the Wnt-enhancesome transcriptional complex (*bcl9, pygo1/2, smarca4a*) (**Fig. 3G**) and downstream Wnt/β-catenin target genes (*tcf7, lef1, jun, fosab*) (**Fig. 3H**), followed the same pattern: all had elevated gene expression with loss of *prdm3* and conversely, all had decreased expression with loss of *prdm16*.

**Figure 3.**
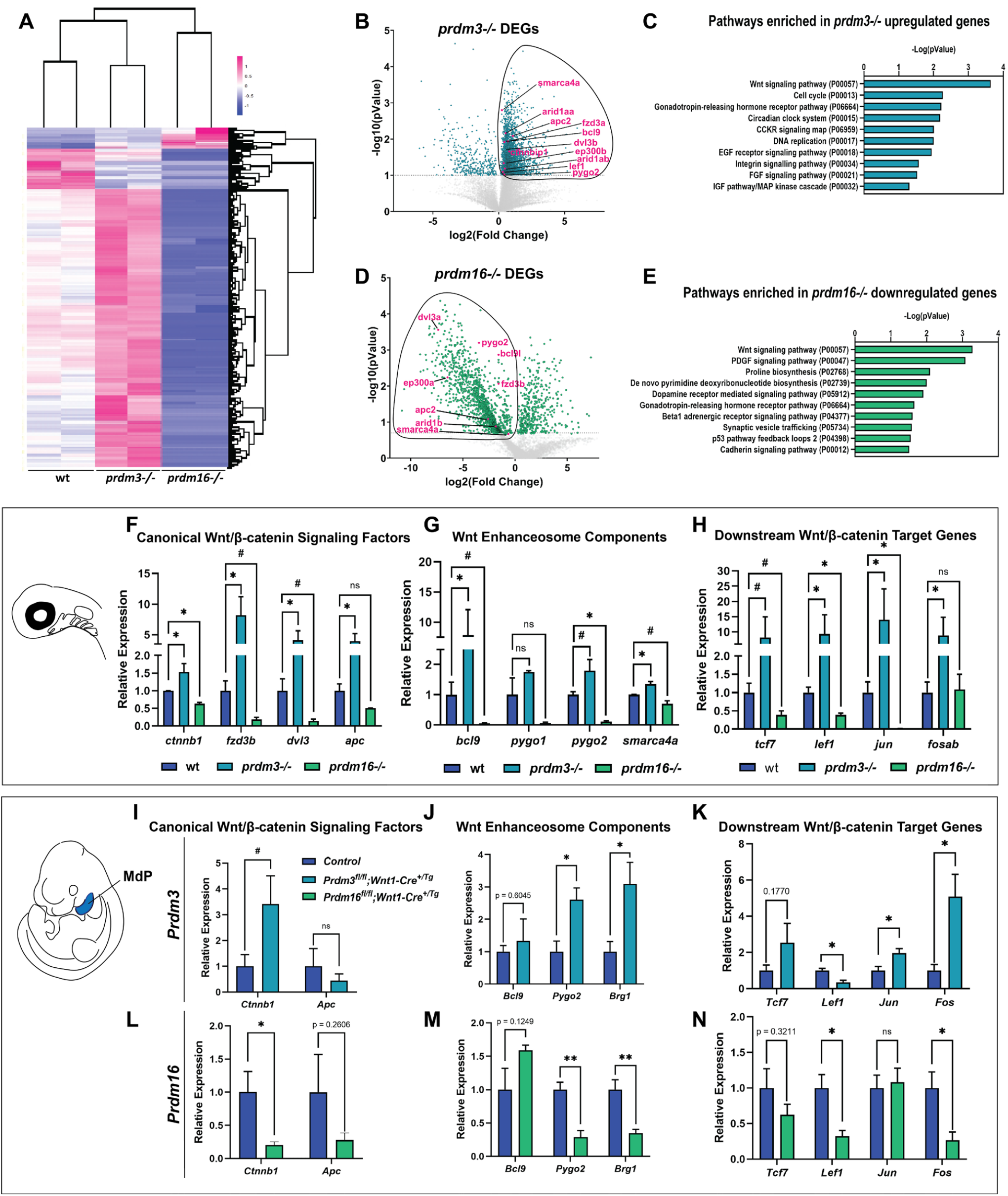
Prdm3 and Prdm16 differentially regulate Wnt/β-catenin signaling components in cranial neural crest cells. (A-E) RNA-sequencing was performed on *sox10:EGFP* FAC-sorted cranial neural crest cells at 48 hpf from wildtype, *prdm3-/-* and *prdm16-/-*. (A) Heatmap showing all significantly differentially expressed genes from *prdm3-/-* and *prdm16-/-* mutants. (B, D) Volcano plots showing the distribution of differentially expressed genes (DEGs) in *prdm3-/-* (B) and *prdm16-/-* (D), revealing strikingly different transcriptomic profiles between the two mutants, many of the same genes are oppositely regulated (upregulated in *prdm3* mutants, downregulated in *prdm16* mutants), including Wnt/β-catenin signaling components (highlighted in magenta). The upregulated genes in *prdm3* mutants and downregulated genes in *prdm16* mutants (circled) were subjected to GO (PANTHER) Pathway Enrichment Analysis (C, E). The Wnt signaling pathway was the most significantly enriched pathway in both gene sets (C, E). (F-H) Validation of RNA-seq transcriptomic changes on RNA isolated from whole heads pooled from 5-7 embryos per genotype. qRT-PCR was performed for selected Wnt/β-catenin signaling component genes including canonical Wnt/β-catenin signaling effectors (*ctnnb1, fzd3b, dvl3, apc*) (F); Wnt enhanceosome transcriptional complex factors (*bcl9, pygo1, pygo2, smarca4a*) (G); and downstream Wnt/β-catenin target genes (*tcf7, lef1, jun, fosab*) (H). (I-N) Validation of zebrafish RNA-seq transcriptomic changes in E11.5 mouse mandibular processes. q-RT-PCR was performed for selected genes including canonical Wnt/β-catenin signaling effectors (*Ctnnb1, Apc*) in *Prdm3* (I) and *Prdm16* (L) mutants; Wnt enhanceosome transcriptional complex factors (*Bcl9, Pygo2, Brg1* (*smarca4a*)) in *Prdm3* (J) and *Prdm16* (M) animals; and downstream Wnt/β-catenin target genes (*Tcf7, Lef1, Jun, Fos*) in *Prdm3* (K) and *Prdm16* (N) mutants. n = 3 for each genotype. MdP, mandibular process; MxP, maxillary process; FL, forelimb. * p ≤ 0.05, # p ≤ 0.1, ns, not significant, Student’s *t* test.

To determine if changes to the expression of Wnt/β-catenin signaling components identified from RNA-seq in zebrafish were also differentially expressed during chondrocyte development in the formation of the mouse Meckel’s cartilage, the mandibular processes of pharyngeal arch 1 at E11.5 control, *Prdm3*^*fl/fl*^*;Wnt1-Cre*^*+/Tg*^, or *Prdm16*^*fl/fl*^*;Wnt1-Cre*^*+/Tg*^ mutant embryos were dissected and the overlying ectoderm removed. RNA was extracted from the mesenchyme of these mandibular facial processes and qRT-PCR was performed for Wnt/β-catenin signaling components (**Fig. 3I-3N**). As with loss of *prdm3* in zebrafish, loss of *Prdm3* in developing murine mandibular facial tissue led to an increase in *Ctnnb1* (**Fig. 3I**) expression, as well as members of the Wnt enhanceosome (*Bcl9, Pygo2, Brg1*) (**Fig. 3J**) and subsequent downstream Wnt/β-catenin target genes (*Tcf7, Jun, Fos*) (**Fig. 3K**). Correspondingly, loss of *Prdm16* in these tissues caused a consistent decrease expression of canonical Wnt/β-catenin signaling factors (*Ctnnb1, Apc*) (**Fig. 3L**), members of the Wnt enhanceosome (*Pygo2, Brg1*) (**Fig. 3M**) and downstream Wnt/β-catenin target genes (*Tcf7, Lef1, Jun, Fos*) (**Fig. 3N**). While there were some differences between expression in some of these genes between species (i.e. decreased *Apc* and *Lef1* expression in *Prdm3* mouse tissues that have elevated expression in zebrafish), the overall the trends are consistent between mice and zebrafish and further suggest that Prdm3 and Prdm16 exert opposing roles in regulating Wnt/β-catenin signaling component gene expression in these populations of cranial NCCs and that this is conserved across tetrapods and ray-finned fishes.

Together, these results suggest Prdm3 and Prdm16 transcriptionally control Wnt/β-catenin signaling components in vertebrates to facilitate proper differentiation and maturation of cranial NCC-derived chondrocytes and further maintain chondrocyte functionality over crucial developmental stages during the formation of the craniofacial skeleton. These proteins may not only affect direct expression of Wnt/β-catenin signaling components, but may also facilitate the coordination of transcriptional activation of Wnt/β-catenin by mediating the expression of the key factors involved in driving nuclear localization of β-catenin and assembling the Wnt transcriptional complex (i.e. *bcl9, pygo1/2, smarca4a, ep300a/b*). Furthermore, while these two PRDMs are very much similar to each other in structure and share similar developmental expression patterns, these results suggest that they may have antagonizing roles: *Prdm3* may normally act as a repressor of gene expression while *Prdm16* may act as a transcriptional activator. The corresponding opposite changes in gene expression affecting the same pathways further suggests that these PRDMs are not only acting opposite of each other but are also functioning at similar gene targets during these stages of chondrocyte development in order to maintain a proper temporal and spatial balance of gene expression and subsequent signaling.

### *prdm3* and *prdm16* control β-catenin stabilization and localization in craniofacial chondrocytes

Given canonical Wnt/β-catenin signaling was the most differentially altered pathway in both *prdm3* and *prdm16*, though in opposing directions (upregulated in *prdm3* mutants, downregulated in *prdm16* mutants), and many of the differentially expressed genes associated with Wnt/β-catenin signaling are involved in the formation of the Wnt/β-catenin transcriptional complex and the retention of nuclear β-catenin, we next assessed the subcellular changes in localization of β-catenin. Whole mount immunofluorescence for nuclear β-catenin (phosphorylated tyrosine residue 489) on wildtype, *prdm3-/-*, or *prdm16-/-* larvae was performed at 75 hpf, the timepoint at which chondrocytes in wildtype larvae have properly oriented, intercalated and have begun to mature. Unlike wildtype chondrocytes, which have low abundance of nuclear β-catenin at this stage (**Fig 4A-4A’’’, 4D**), loss of *prdm3-/-* caused a significant increase in the presence of nuclear β-catenin (**Fig. 4B-4B’’’, 4D**). Conversely, loss of *prdm16-/-* caused a dramatic reduction in the accumulation of β-catenin in the nucleus to levels significantly below that of wildtype chondrocytes (**Fig. 4C-4C’’’, 4D**). This was quantified by counting the number of phosphorylated Y-489 β-catenin puncta per nuclei (**Fig. 4D**).

**Figure 4.**
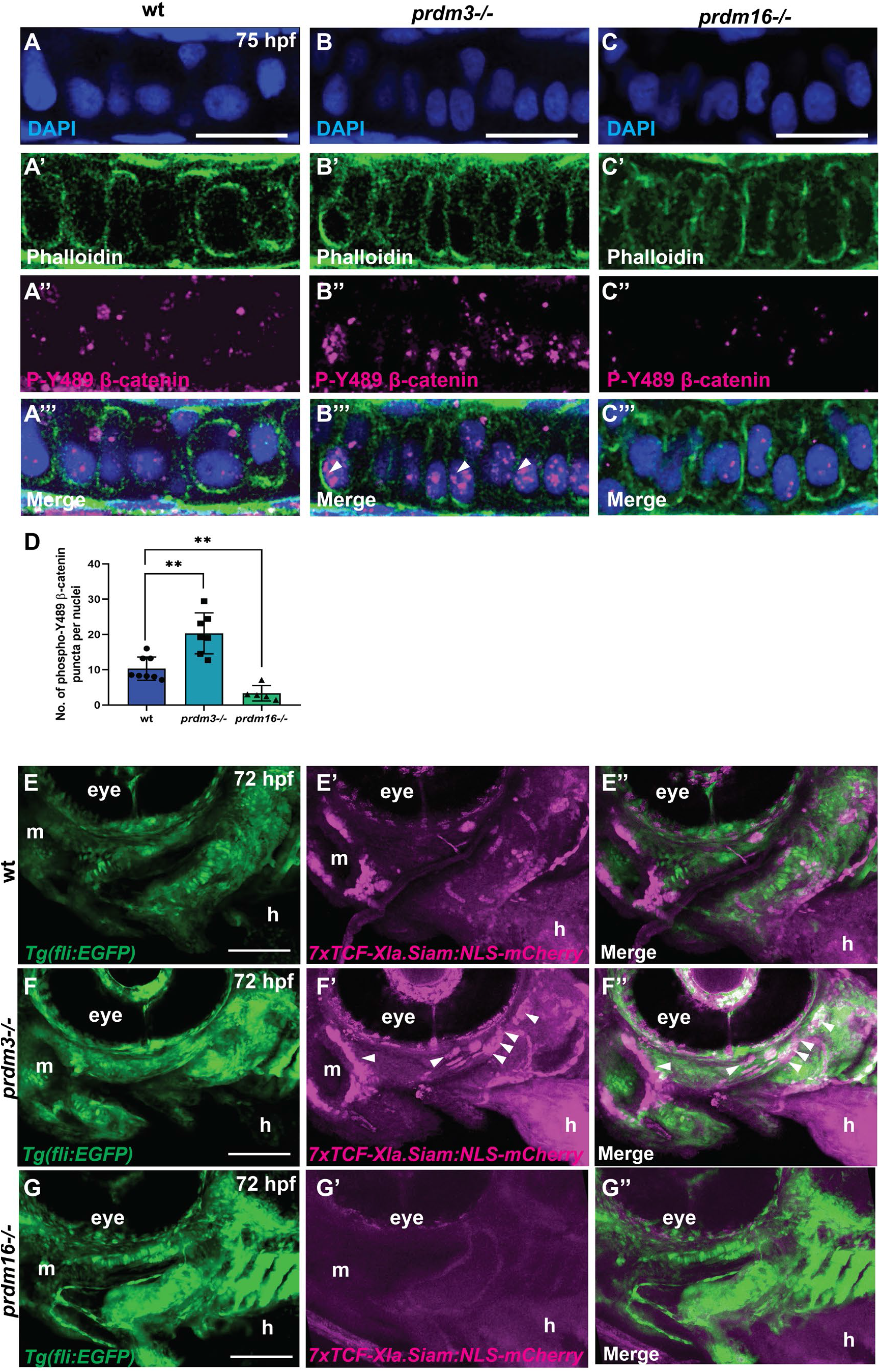
Loss of *prdm3* and *prdm16* alters nuclear accumulation of β-catenin and Wnt responsive cells during craniofacial chondrocyte development in zebrafish. (A-D) Wildtype, *prdm3-/-* and *prdm16-/-* embryos were collected at 75 hpf and whole mount immunofluorescence was performed for phosphorylated tyrosine 489 residue on β-catenin, indicative of nuclear β-catenin (magenta) (A’’-C’’) and counterstained with phalloidin (A’-C’) to label F-actin filaments and DAPI (A-C). Shown are high magnification lateral images of the developing palatoquadrate. Increased nuclear β-catenin was observed in *prdm3-/-* (white arrowheads in B’’’), which was significantly reduced in *prdm16-/-* (C’’’). (D) Quantification of the number of phosphorylated tyrosine 489 puncta across 10 nuclei per individual. Shown are the averages from at least 5 individuals per genotype. Scale bar, 50 µm. (E-G’’) *prdm3-/-* and *prdm16-/-* mutant lines were crossed into the Wnt reporter line, *Tg(7xTCF-Xla*.*Siam:NLS-mCherry)* to assess Wnt responsive cells and *Tg(fli:EGFP)* which labels the developing pharyngeal arch derivatives and vasculature. Shown are representative lateral-ventral views of 75 hpf wildtype (wt) (E-E’’), *prdm3-/-* (F-F’’) and *prdm16-/-* (G-G’’). Increased Wnt responsive cells were identified in the pharyngeal arch tissues of *prdm3-/-* (F-F’’) (white arrowheads), with a dramatic decrease in Wnt responsive cells in *prdm16-/-* (G-G’’) mutants compared to wildtype controls (E-E’’). Scale bar, 100 µm. m, mouth; h, heart. ** p ≤ 0.005, Student’s *t* test.

To further understand canonical Wnt/β-catenin signaling in chondrocyte development in the craniofacial skeleton with loss of *prdm3* and *prdm16, prdm3-/-* and *prdm16-/-* zebrafish mutants were crossed into the Wnt reporter line, *Tg(7xTCF-Xla*.*Sia:NLS-mCherry)*^*ia5Tg*^ [77] in combination with the *Tg(fli1:EGFP)* [78] line to visualize Wnt responsive cells in the pharyngeal arches and developing craniofacial cartilage elements (**Fig. 4E-4G)**. At 72 hpf, *prdm3* mutants developed a dramatic increase in Wnt responsive cells not only in the pharyngeal arch/craniofacial elements, but in other developing structures and tissues of the larvae, including the heart, otic vesicle and perichondrium (**Fig. 4F-4F’’**). Conversely, little to no Wnt responsive cells were present in *prdm16-/-* mutant larvae (**Fig. 4G-4G’’**). Altogether, these results validate the transcriptional changes observed in both *prdm3* and *prdm16* mutants and correlate with differences in expression of factors (i.e. *bcl9)*, involved in shuttling and retaining β-catenin to the nucleus and driving transcriptional activity of downstream target genes. Abnormal accumulation of nuclear β-catenin in *prdm3* mutant chondrocytes at later stages of development further explains the chondrocyte polarity, orientation, intercalation and differentiation defects, as high sustained β-catenin signaling is inhibitory to cartilage differentiation [24, 25, 31, 79]. On the other hand, while *prdm16* mutants have these same chondrocyte morphological and differentiation defects, affecting the same signaling pathways, but instead with a dramatic reduction of canonical Wnt/β-catenin signaling and absence of low-level maintenance of nuclear β-catenin that would otherwise help facilitate chondrocyte differentiation and maturation.

### *prdm3* and *prdm16* alter chromatin accessibility at *cis*-regulatory regions of Wnt/β-catenin specific targets in cranial NCCs

Previous work has implicated both Prdm3 and Prdm16 in regulation of gene expression through methylation of lysine residues on histone H3— H3K9, a mark of repression and H3K4, a mark associated with gene activation. Loss of *prdm3* and *prdm16* significantly reduces H3K9me3 and H3K4me3 in zebrafish [72], suggesting global changes to the chromatin landscape, that ultimately affect both gene activation and gene repression. To better understand how Prdm3 and Prdm16 are epigenetically regulating gene expression, specifically of canonical Wnt/β-catenin pathway components, assays for transposase-accessible chromatin paired with sequencing (ATAC-seq) were performed on FAC-sorted *sox10:EGFP* cranial NCCs isolated from pooled wildtype, *prdm3-/-* and *prdm16-/-* embryos at 48 hpf (**Fig. 5**). Loss of *prdm3* led to an increase in open chromatin with 89.6% open peaks compared to 10.4% in wildtype (**Fig. 5A**). Contrary to this, loss of *prdm16* caused an overall decrease in open chromatin with only 34.3% open peaks compared to 65.7% in wildtype (**Fig. 5C**). The increased areas of open chromatin in *prdm3-/-* were associated with promoters, within regions 2.5 kb upstream of transcriptional start sites (TSS) across the genome (**Fig. 5B, 5F**) compared to wildtype (**5B, 5E**). There were no differences in open chromatin at other genomic regions (exons, introns, intergenic, 5’UTR, 3’UTR, TTS) (**Fig. 5B**). The decrease in open chromatin in *prdm16-/-* did not account for any changes in the genomic distribution of peaks (**Fig. 5D**) and areas of accessibility were decreased immediately upstream and downstream of TSSs compared to wildtype (**Fig. 5E, 5G**).

**Figure 5.**
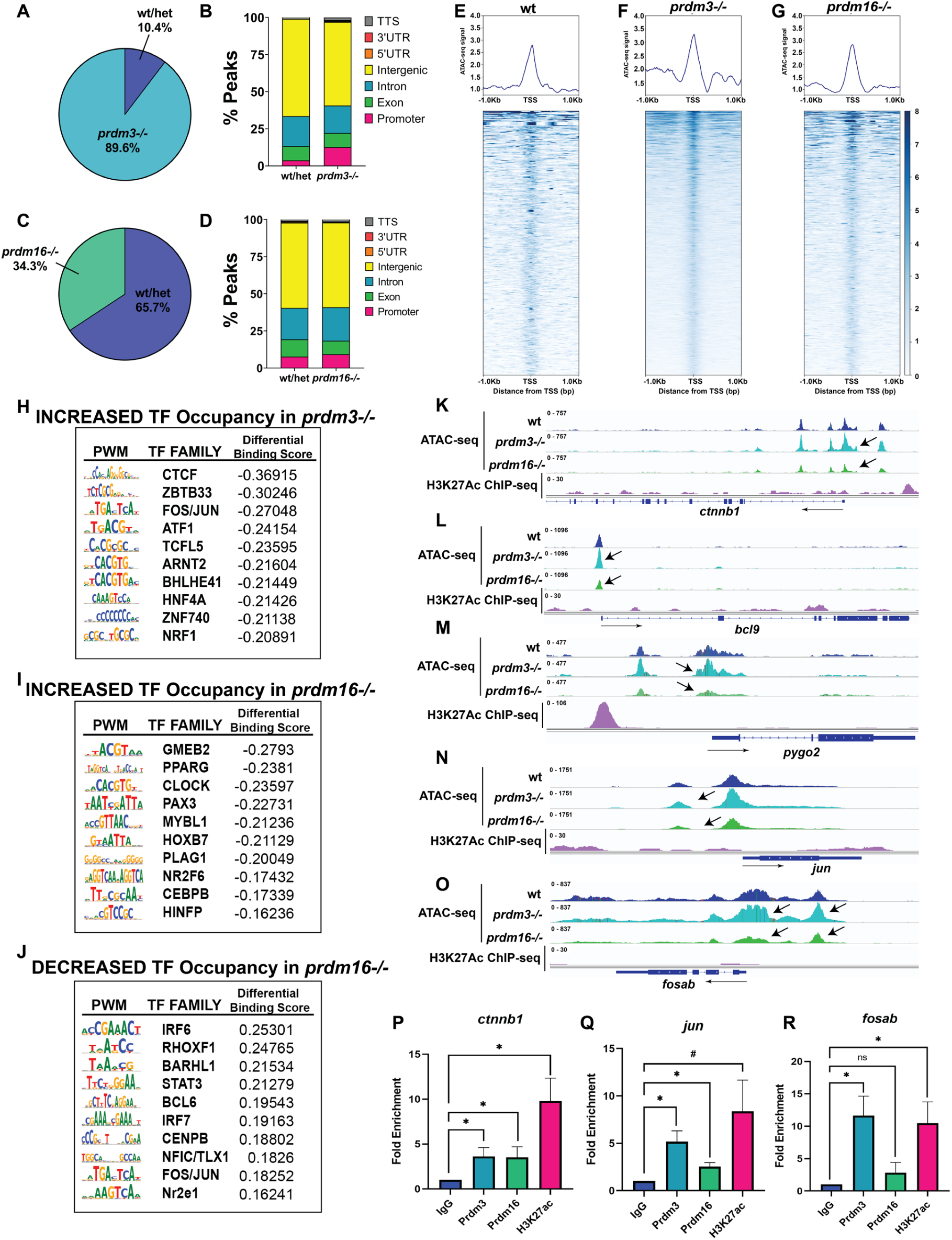
Loss of *prdm3* and *prdm16* alters chromatin accessibility at cis-regulatory regions of Wnt/β-catenin signaling components in cranial neural crest cells. (A-O) Assay for Transposase Accessible Chromatin (ATAC) paired with sequencing was performed on *sox10:EGFP* FAC-sorted cells from wildtype, *prdm3-/-* and *prdm16-/-* at 48 hpf. (A, C) Percentage of significant peaks identified in *prdm3-/-* (A) and *prdm16-/-* (C) compared to control. (B, D) Annotation of the location of peaks in *prdm3-/-* (B) shows more peaks are associated with promoter regions compared to controls, with no change in the location of peak distributions in *prdm16-/-* (D). (E-G) Coverage heatmaps depicting the differences in peak coverage (indicative of open chromatin) 1kb upstream of transcriptional start sites (TSSs) across the genome, with an increase of open chromatin 1kb upstream of TSS in *prdm3-/-* (F) and decreased chromatin accessibility in *prdm16-/-* (G) compared to controls (E). (H-J) Transcription factor Occupancy prediction By Investigation of ATAC-seq Signal (TOBIAS) was performed on *prdm3* and *prdm16* mutant ATAC datasets to predict which transcription factors are bound in with loss of *prdm3* or *prdm16*. Shown are the top 10 transcription factors occupying promoter regions in *prdm3-/-* (H) and *prdm16-/-*, p values ≤ E-20. (I). Because the overwhelming majority of genes were downregulated in *prdm16-/-* which corresponded to an overall decrease in accessible chromatin, transcription factors that had decreased occupancy were also assessed in *prdm16-/-* (J). Each table shows the PWM (position weight matrix) for each transcription factor motif, the family that transcription factor belongs to, the differential binding score produced from TOBIAS, and the p-value. (K-O) Tracks showing the distribution of peaks (chromatin accessibility) (same scale range across genotypes for each gene), compared to previously published H3K27ac ChIP-seq datasets [88] from 48 hpf whole embryos, at specific Wnt/β-catenin signaling component target genes that were differentially-expressed in *prdm3* and *prdm16* mutants including *ctnnb1* (K), *bcl9* (L), *pygo2* (M), *jun* (N), and *fosab* (O). (P-R) Cleavage Under Targets and Release Using Nuclease (CUT&RUN) paired with qRT-PCR was performed on 48 hpf whole embryos with antibodies directed toward Prdm3, Prdm16, and H3K27ac. Enrichment for Prdm3, Prdm16, and H3K27Ac was assessed with primers designed to flank putative binding sites of Prdm3 and Prdm16 near the promoter regions (defined as within 1000 bp of the TSS) of these genes, including: *ctnnb1* (P), *jun* (Q), and *fosab* (R). * p ≤ 0.05, ns, not significant, Student’s *t* test.

To predict the enrichment of differentially bound transcription factors in *prdm3-/-* and *prdm16-/-* cNCCs, we utilized the computational footprinting pipline TOBIAS (Transcription factor Occupancy prediction by Investigation of ATAC-seq Signal) [80]. In *prdm3-/-* mutants, the top 10 transcription factor families with increased binding were associated with regulating genes necessary for cell growth, differentiation and developmental processes (**Fig. 5H, See also Supplemental Table 2 and Supplemental Table 3**). Several of these transcription factors (ZBTB33, CTCF) can also recruit protein complexes such as N-CoR (ZBTB33) or other remodeling factors (histone deacetylases, histone methyltransferases) to modulate chromatin architecture and form complexes for DNA looping to drive gene expression through enhancers (**Fig. 5H**). Interestingly, one of the top transcription factor families differentially bound in *prdm3-/-* mutant cNCCs was Fos/Jun, a top transcriptional target of canonical Wnt/β-catenin signaling. *fos* and *jun* were also both differentially expressed at the transcriptional level in both *prdm3* and *prdm16* mutants (**Fig. 3**).

The top 10 transcription factors bound in *prdm16-/-* cranial NCCs were strikingly different from those identified in *prdm3* mutants and instead included several factors that Prdm16 is already known to interact with including PPARG and CEBPB to regulate brown fat adipogenesis [63, 64] (**Fig. 5I, See also Supplemental Table 4**). PPARG and Wnt/β-catenin can antagonistically influence expression of each other—activated PPARG decreases Wnt/β-catenin and elevated Wnt/β-catenin inhibits PPARG [81-83]. This pattern of Wnt/β-catenin expression and occupancy of PPARG correlates with the decreased Wnt/β-catenin observed in *prdm16* mutants. Other transcription factors important for differentiation (PAX3) and patterning and cell adhesions (HOXB7) in development had increased occupancy in *prdm16* mutants. Because most genes were downregulated and associated with less accessible chromatin with loss of *prdm16*, the transcription factors that were significantly unbound or lost in these mutants were also assessed (**Fig. 5J, See also Supplemental Table 5**). The top transcription factor with significantly decreased binding with loss of *prdm16* was IRF6. IRF6 has important functions in regulating cell-cell adhesions and plays a crucial role in murine palatal shelf fusion [84-87]. Other transcription factors lost in *prdm16* mutants included Wnt pathway target, Bcl6. Fos/Jun binding was also decreased in *prdm16-/-* cranial NCCs which further correlates to the opposite gain of Fos/Jun occupancy with loss of *prdm3*.

As canonical Wnt/β-catenin signaling was the most differentially altered pathway in both *prdm3* and *prdm16* mutants, changes in chromatin accessibility were assessed at different Wnt/β-catenin signaling components and target genes (**Fig. 5K-5O**). Consistent with our transcriptomic data, loss of *prdm3* led to increased open chromatin at canonical Wnt/β-catenin components, including *ctnnb1*(**Fig. 5K**), *blc9* (**Fig. 5L**), *pygo2* (**Fig. 5M**) as well as downstream Wnt/β-catenin target *jun* (**Fig. 5N**) and binding partner *fos* (**Fig. 5O**). Conversely, chromatin accessibility of these Wnt/β-catenin targets was dramatically decreased in *prdm16-/-* (**Fig. 5K-5O**). Areas of open chromatin aligned with areas of active gene expression as assessed with previously published H3K27Ac ChIP-seq data from whole zebrafish embryos at 48 hpf [88]. To determine if *prdm3* and *prdm16* can directly bind to promoter regions of these canonical Wnt/β-catenin signaling component genes and mediate their expression, cleavage under targets and release using nuclease paired with quantitative real-time PCR (CUT&RUN-qRT-PCR) was performed on whole zebrafish larvae at 48 hpf. Significant enrichment of Prdm3 and Prdm16 abundance was identified at putative binding sites for Prdm3 and Prdm16 near promoter regions of *ctnnb1* and downstream Wnt/β-catenin target gene, *jun*. Only Prdm3 had significant enrichment at *fosab*, suggesting differences in target genes between Prdm3 and Prdm16. Decreased expression of *fosab* in *prdm16* mutants with no changes in Prdm16 enrichment at this gene suggests this is not direct target of Prdm16 and instead is indirectly differentially expressed due to changes in expression upstream Wnt/β-catenin transcriptional regulatory components. There was no enrichment of Prdm3 or Prdm16 at promoter regions of *bcl9* or *pygo2* (**Supplemental Fig. 4C-4D**). While this result suggests these factors are not direct transcriptional targets Prdm3 or Prdm16, this does not exclude their possible protein-protein interaction and subsequent influence on Wnt/β-catenin transcriptional activity. Taken together, these results suggest Prdm3 and Prdm16 may not only indirectly influence transcription of Wnt/β-catenin signaling components by changing chromatin accessibility near promoter regions of these genes but may also affect expression of these Wnt/β-catenin genes by binding those targets directly. These results further emphasize opposing roles of *prdm3* and *prdm16* in facilitating a balance of canonical Wnt/β-catenin signaling during cNCC-derived chondrocyte differentiation in the craniofacial skeleton. While Prdm3 and Prdm16 enrichment is detected at some of these Wnt/β-catenin signaling components, the changes of gene expression of these factors are completely opposite. These results provide further evidence that in this cellular context under normal developmental conditions, *prdm3* may act as a transcriptional repressor while *prdm16* serves as an activator of gene expression. Additionally, loss of either alters the chromatin landscape by changing accessibility in opposing directions, in coordination with changes in gene expression, which disrupts downstream transcriptional networks.

### Pharmacological manipulation of canonical Wnt/β-catenin signaling in *prdm3-/-* and *prdm16-/-* zebrafish mutants partially restores chondrocyte stacking defects

Our genomic data suggest that Prdm3 and Prdm16 are regulating Wnt activity in opposing ways. We hypothesize that these transcription factors may be acting to titer Wnt/β-catenin transcriptional activity during chondrocyte differentiation. To test this, we next sought to rescue cartilage phenotypes by pharmacologically manipulating Wnt signaling in both zebrafish mutants. To rescue the effects of elevated Wnt/β-catenin signaling and subsequent altered chondrocyte cell organization and adherens junctions observed in *prdm3* mutants, wildtype and *prdm3-/-* embryos were treated with either DMSO or 0.75 μM of the Wnt inhibitor, IWR-1, from 24 hpf to 48 hpf. IWR-1 blocks Wnt-induced β-catenin accumulation by stabilizing the Axin2 destruction complex [89]. At 6 dpf, control and treated embryos were collected and stained with Alcian blue and alizarin red to assess cartilage and bone (**Fig. 6A-6A’’’**). The chondrocytes in vehicle treated *prdm3* mutants were highly disorganized (**Fig 6A’**). IWR-1 treatment significantly restored chondrocyte organization and intercalation within these mutant cartilage structures (**Fig. 6A’’’**), as quantified by measuring the angle between adjacent chondrocytes (**Fig. 6B**).

**Figure 6.**
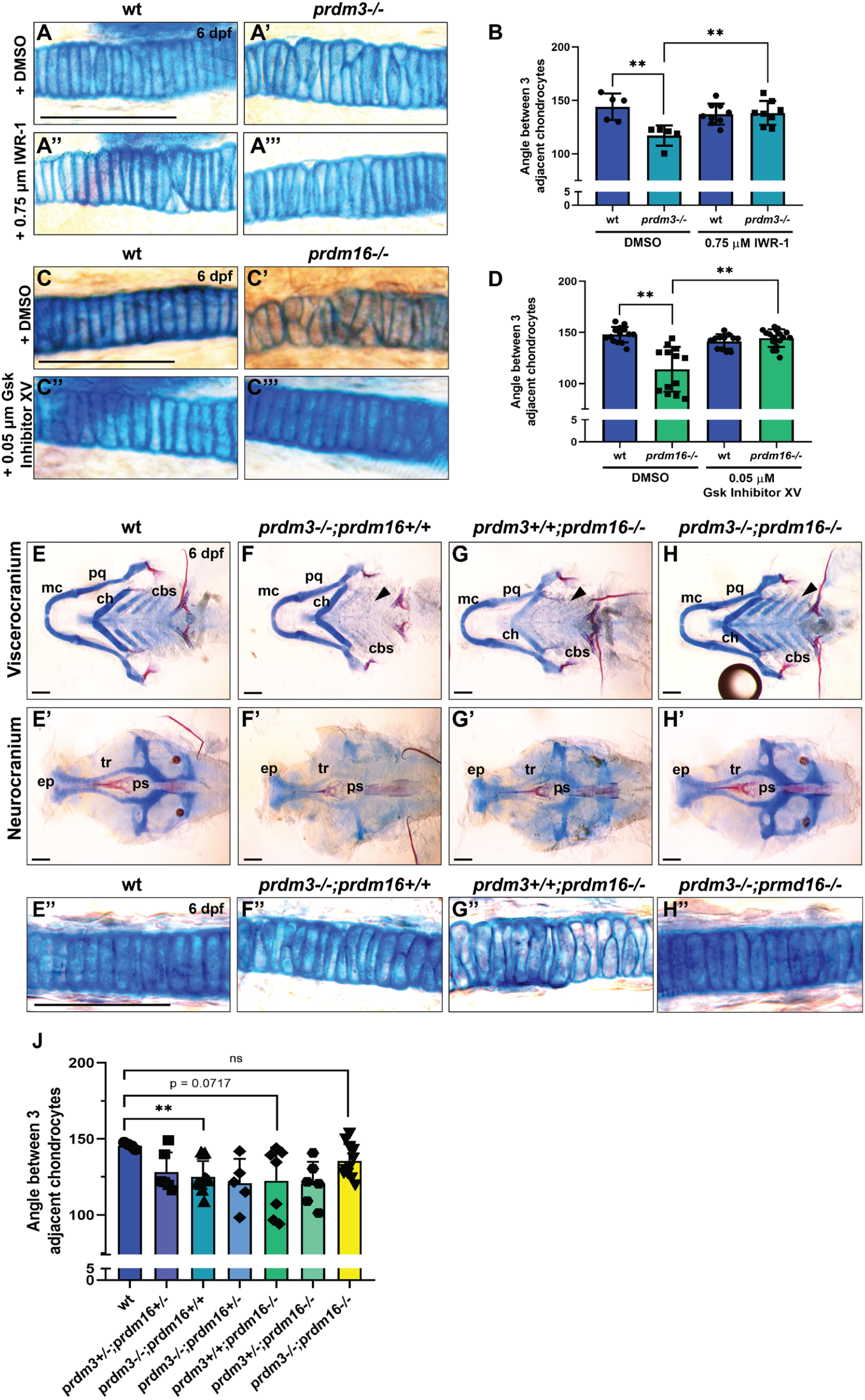
Chondrocyte stacking defects with loss of *prdm3* and *prdm16* can be rescued by pharmacologically manipulating Wnt/β-catenin activity or genetically with *prdm3-/-;prdm16-/-* double mutants. (A-B) Wildtype (wt) or *prdm3-/-* embryos were treated with suboptimal doses of 0.75 µM IWR-1 or DMSO (vehicle control) from 24 hpf to 48 hpf. Inhibitor and vehicle treated larvae were collected at 6 dpf and stained with Alcian blue or Alizarin red. (A-A’’’) High magnification of chondrocytes in cartilage elements showed abnormal chondrocyte stacking with vehicle treated *prdm3-/-* mutants (A’) that were rescued with IWR-1 (Wnt pathway inhibition) treatment (A’’’). (B) Quantification of chondrocyte stacking and organization. Measurements were collected from at least 5 individuals per genotype per treatment group. Scale bar, 100 µm. (C-D) Wildtype (wt) or *prdm16-/-* embryos were treated with suboptimal doses of 0.05 µM Gsk Inhibitor XV (Wnt activator) or DMSO (vehicle control) from 24 hpf to 48 hpf. Larvae were collected at 6 dpf and stained with Alcian blue and Alizarin red. (C-C’’’) High magnification of chondrocytes within cartilage elements showed altered chondrocyte stacking in vehicle treated *prdm16-/-* mutants (C’) compared to controls (C) that were rescued with Gsk Inhibitor XV (Wnt pathway activation) treatment (C’’’). (D) Quantification of chondrocyte organization. Measurements were collected from at least 10 individuals per genotype per treatment group. Scale bar, 100 µm. (E-H’’) *prdm3+/-;prdm16+/-* heterozygous fish were generated and intercrossed. Larvae were collected at 6 dpf and stained with Alcian blue and Alizarin red. (E-H, E’-H’) Shown are dissected and mounted viscerocranium (E-H) and neurocranium (E’-H’) from wildtype (E, E’), single *prdm3* mutants (F, F’), single *prdm16* mutants (G, G’) and *prdm3-/-;prdm16-/-* double mutants (H, H’). (E’’-H’’) High magnification of chondrocytes in cartilage elements of wildtype (E’’), *prdm3* single mutants (F’’), *prdm16* single mutants (G’’) and *prdm3-/-;prdm16-/-* double mutants (H’’). (J) Quantification of the angle between adjacent chondrocytes across wildtype, *prdm3-/-, prdm16-/-* and *prdm3-/-;prdm16-/-* double mutants, as well as all other combinatorial mutants indicated. cbs, ceratobranchials; ch, ceratohyal; ep, ethmoid plate; mc, Meckel’s cartilage; pq, palatoquadrate; ps, parasphenoid; tr, trabeculae. Scale bar, 100 µm. ** p ≤ 0.005, ns, not significant, Student’s *t* test.

To mitigate the effects of decreased Wnt/β-catenin signaling and the same subsequent chondrocyte polarity and stacking defects in *prdm16-/-* larvae, wildtype and *prdm16-/-* embryos were treated with either DMSO or 0.05 µM of a Wnt/β-catenin activator, GSK-3 Inhibitor XV, from 24 hpf to 48 hpf. GSK Inhibitor XV blocks Gsk3-dependent phosphorylation of β-catenin allowing for β-catenin stabilization. Larvae were collected at 6 dpf and Alcian blue and Alizarin red stained (**Fig. 6C-6C’’’**). While chondrocytes of *prdm16-/-* vehicle treated larvae exhibited stacking and intercalation defects (**Fig. 6C’**), Wnt/β-catenin activation (via Gsk3 inhibition) completely rescued chondrocyte cellular defects (**Fig. 6C’’’**). This was quantified by measuring the angle between adjacent chondrocytes (**Fig. 6D**). GSK Inhibition (Wnt pathway activation) also completely rescued the overall cartilage phenotypes in the craniofacial skeleton of *prdm16-/-* mutants, including restoration of posterior ceratobranchial cartilages (**Supplemental Fig. 5A**). Together, these results suggest loss of *prdm3* and *prdm16* compromises chondrocyte intercalation and maturation during craniofacial development and these defects can be restored by rebalancing Wnt/β-catenin signaling pharmacologically in each mutant.

### Combined genetic loss of both *prdm3* and *prdm16* rescues chondrocyte stacking defects

While our genomic data suggest that Prdm3 and Prdm16 are regulating Wnt/β-catenin activity in opposing directions, we found that single loss of *prdm3* or *prdm16* causes similar cartilage phenotypes: defects in chondrocyte orientation, polarity, intercalation and downstream maturation. To determine if genetic loss of both *prdm3* and *prdm16* would rescue or exacerbate the craniofacial cartilage phenotypes observed in the single mutants, double homozygous *prdm3-/-;prdm16-/-* mutants (and all other allelic combinations) were generated from *prdm3+/- ;prdm16+/-* heterozygous intercrosses (**Fig. 6E-6H**). Surprisingly, loss of both *prdm3-/-* and *prdm16-/-* rescued craniofacial phenotypes observed in single mutant embryos. *prdm3-/- ;prdm16-/-* larvae had normal posterior arch cartilage structures, and partial restoration of the hypoplasia observed in other cartilage elements (Meckel’s cartilages, trabeculae, ethmoid plate) (**Fig. 6H-6H’**). *prdm3-/-;prdm16-/-* double mutants also had a near complete rescue of chondrocyte stacking and intercalation defects that were observed in single mutants (**Fig. 6H’’, 6J**). Combinatorial loss of single mutants with heterozygous loss of the other corresponding alleles did not rescue overall cartilage phenotypes or cellular stacking defects suggesting the rescue is dependent on complete loss of both paralogs (**Fig. 6J, Supplemental Fig. 5B-5C**). Taken together, these results emphasize the functional roles of *prdm3* and *prdm16* in working not only together, but also independent of each other to facilitate a balance of gene regulatory networks and signaling modules during craniofacial cartilage development such as canonical Wnt/β-catenin signaling.

## Discussion

This study defines the mechanistic functions of PRDM3 and PRDM16 in chondrocyte differentiation during the development of the craniofacial complex in zebrafish and mice (**Fig. 7**). We show that loss of Prdm3 and Prdm16 impairs proper chondrocyte orientation, polarity and intercalation. Transcriptomic analysis and assessment of the chromatin landscape revealed strikingly different epigenetic and transcriptomic profiles. Loss of *prdm3* led to an overwhelming increase in open chromatin and a subsequent increase in gene expression. In contrast, loss of *prdm16* caused a dramatic decrease in chromatin accessibility and a significant downregulation of gene expression. With these completely opposite effects on gene expression and chromatin architecture, both *prdm3* and *prdm16* modulate the same signaling pathways, specifically canonical Wnt/β-catenin signaling. We demonstrate we can rescue chondrocyte stacking and intercalation defects with loss of *prdm3* and *prdm16* by both pharmacologically restoring Wnt/β-catenin signaling levels in each mutant and genetically with loss of both *prdm3* and *prdm16* in *prdm3-/-;prdm16-/-* double mutants. Lastly, we show conservation of these mechanisms during mammalian craniofacial development with changes to chondrocyte organization in the developing mandible and altered expression of Wnt/β-catenin signaling components in the mandibular facial prominences in *Prdm3* and *Prdm16* mutants.

**Figure 7.**
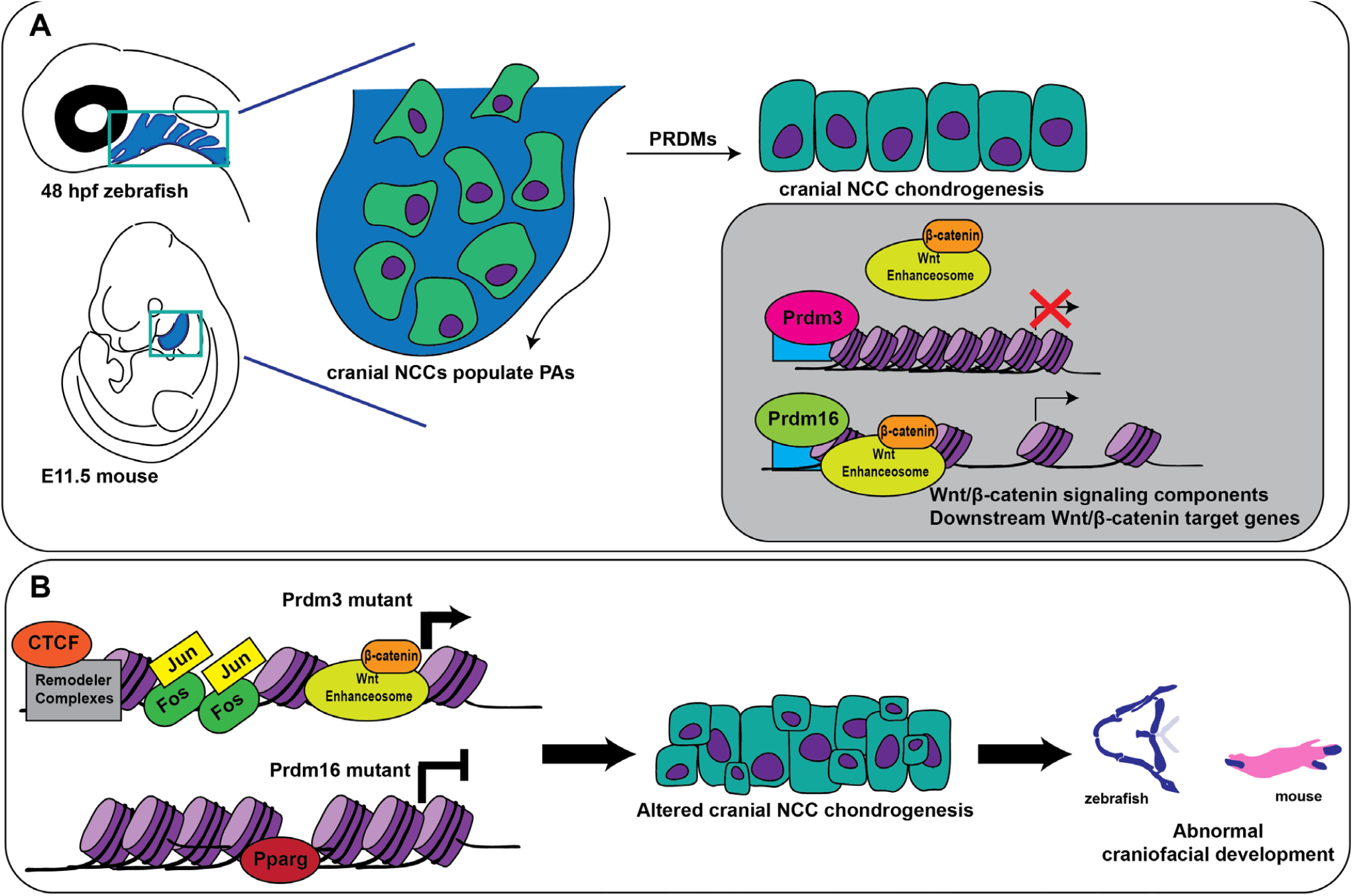
Prdm3 and Prdm16 control craniofacial chondrogenesis by balancing upstream Wnt/β-catenin transcriptional activity. (A) In vertebrates, Prdm3 and Prdm16 facilitate cranial NCC chondrocyte differentiation and maturation by balancing the temporal and spatial upstream regulation of Wnt/β-catenin transcriptional activity, where Prdm3 acts a repressor of gene expression and Prdm16 acts as an activator of similar gene targets, particularly Wnt/β-catenin signaling components and downstream Wnt/β-catenin target genes. (B) Loss of Prdm3 leads to enhanced gene expression and increased occupancy of CTCF, Jun/Fos while loss of Prdm16 causes a dramatic decrease in gene expression and increased occupancy of Pparg, among others. In both cases, Wnt/β-catenin signaling is abrogated leading to altered cranial neural crest chondrocyte differentiation and maturation which ultimately leads to abnormal development of craniofacial structures. PA, pharyngeal arches; NCC, neural crest cells.

We have identified novel functional roles for Prdm3 and Prdm16 in cranial NCC-derived chondrocyte differentiation during craniofacial development. These Prdms have long been thought to act redundantly of each other, given their highly similar amino acid sequence homology and developmental expression patterns. Contrary to this paradigm, we hypothesize under normal developmental conditions in the context of cranial NCCs, *Prdm3* functions as a traditional transcriptional repressor while *Prdm16* acts as a non-traditional activator of gene expression to regulate cranial neural crest chondrocyte differentiation. Intriguingly, both of these mutants, despite their opposite transcriptomic profiles and chromatin architecture have the same cartilage phenotypes: impaired chondrocyte orientation, polarity, intercalation and growth that subsequently cause abnormal chondrocyte differentiation and maturation. While little is known about the regulatory factors that control expression of Wnt/β-catenin signaling components, we hypothesize that these two chromatin modifiers balance activation and repression of gene regulatory networks and signaling modules, particularly canonical Wnt/β-catenin signaling in order to facilitate proper spatial and temporal activation and repression of this pathway during craniofacial chondrogenesis. As a result, combined loss of both *prdm3* and *prdm16* in *prdm3-/-;prdm16-/-* instead of exacerbating the cartilage phenotypes, rescues not only the cellular stacking defects but the overall craniofacial skeletal malformations. In a Goldilocks like effect, there must be a basal level of Wnt/β-catenin activity during these stages of chondrocyte differentiation as too much or too little activity can have consequential effects on the proper maturation and differentiation of these cells. This result further suggests that there is another level of regulation whereby these highly similar paralogs not only function together, but that each has their own divergent and independent role in facilitating proper chondrocyte and bone differentiation. Interestingly, Prdm3 and Prdm16 result from a vertebrate specific duplication that occurred at the Gnathostomata ancestor [90]. As such, independent functions of Prdm3 and Prdm16, at least in the case of chondrocyte maturation and bone differentiation in craniofacial development, may have coincided with their duplication event which correlates with the evolution of jawed vertebrates.

We have identified a gene regulatory network centered on facilitating temporal and spatial canonical Wnt/β-catenin signaling during cartilage development. It has been long established that β-catenin has a multitude of roles in chondrocytes from signaling at cell-cell adhesions to driving gene expression changes over the course of differentiation. Furthermore, β-catenin, while important in early pre-chondrogenic condensations, needs to be waned to allow for differentiation to continue and active again at late stages of terminal hypertrophic chondrocytes to promote osteoblast differentiation. Here we have identified upstream functional roles for Prdm3 and Prdm16 in facilitating the temporal and spatial regulation of canonical Wnt/β-catenin signaling necessary for proper chondrocyte differentiation. We hypothesize there are several mechanisms both direct and indirect that may be facilitating this temporal regulation. Among the differentially expressed Wnt/β-catenin signaling components, the intermediary protein that facilitates β-catenin nuclear localization and subsequent downstream transcription of target genes, *bcl9*, and it’s binding partner, *pygo1/2*, were significantly altered in both *prdm3* (upregulated) and *prdm16* (downregulated) mutants. Previous work has shown *bcl9* zebrafish mutants have cranial NCC defects including craniofacial cartilage phenotypes [91]. It would be interesting to determine how *prdm3* and *prdm16* regulate *bcl9* to control β-catenin transcriptional activation or whether Prdm3 and Prdm16 participate in the Wnt-enhanceosome transcriptional complex through protein interactions with members of the complex, including *bcl9* or *pygo1/2* to influence Wnt/β-catenin transcriptional activity.

While we have focused only on canonical Wnt/β-catenin signaling, and its impacts on mediating cell-cell adhesions and downstream transcription of targets during chondrocyte differentiation, it is possible non-canonical Wnt/Planar Cell Polarity (PCP) signaling is also active. Recent studies have suggested crosstalk between the two pathways coordinates developmental processes [92]. In alignment with these studies, we did observe significant changes in expression of several Wnt/PCP genes (*vangl2, scrib, fat*) as well as the transcription factors *jun/fos*. While *jun* and *fos* are also a direct canonical Wnt/β-catenin target genes, *jun* and *fos* are also activated in response to non-canonical Wnt/PCP through JNK signaling. Because both *prdm3* and *prdm16* mutants share similar cartilage phenotypes with those observed in PCP zebrafish mutants, we speculate that Prdm3 and Prdm16 could also control crosstalk between both canonical and non-Wnt/β-catenin signaling activity during chondrocyte differentiation. Interestingly, Prdm3 has been shown to directly interact with c-Fos/Jun *in vitro* [93]. We hypothesize that one such mechanism governing canonical and non-canonical Wnt crosstalk could be through Prdm3’s direct interaction with Fos and Jun during cranial NCC cartilage development.

We have identified putative cis-regulatory regions of Wnt/β-catenin target genes that are regulated by Prdm3 and Prdm16. Previous work has implicated Prdm16 in epigenetically regulating enhancer regions to drive gene regulatory networks. Identification of shared and different enhancers regulated by Prdm3 and Prdm16 will facilitate our understanding of how these factors control specific gene regulatory networks during craniofacial development. In return, it could be possible that Prdm3 and Prdm16 are downstream Wnt/β-catenin transcriptional targets which would form a regulatory circuit that could facilitate the timely gradient of Wnt/β-catenin necessary during chondrocyte differentiation and maturation during craniofacial development.

Finally, we have previously identified conserved roles for these factors across species with similarities between the development of posterior arch derivatives including the mammalian middle ear and the telost jaw structures, as well as the mammalian laryngeal cartilages and the telost gill arch structures [72]. Here, we further define the conserved functions of Prdm3 and Prdm16 across vertebrates and their roles in mediating chondrocyte development by controlling expression of Wnt/β-catenin signaling components in craniofacial skeletal formation. We show loss of *Prdm3* and *Prdm16* in the neural crest lineage disrupts chondrocyte organization and maturation within the developing Meckel’s cartilage in the mandible as a result of dysregulated Wnt/β-catenin signaling during early chondrogenesis in the mandibular facial processes. We hypothesize that these changes in chondrocyte maturation could influence the mineralization and development of the mandible, which could explain the anterior mandibular hypoplasia seen in these animals. We hypothesize this mandibular hypoplasia could lead to misplacement of the tongue which, alongside other factors could partially explain the development of secondary cleft palate, observed in the *Prdm16* mutants. It will be interesting to further understand how Prdm3 and Prdm16 regulate this potential coupling mechanism between chondrocyte maturation and subsequent bone formation in the developing mandible. Furthermore, it will be interesting to understand the mammalian genetic interaction between *Prdm3* and *Prdm16* and the subsequent effects on Wnt/β-catenin transcriptional activity during cartilage development in the craniofacial skeleton.

In summary, we have identified novel roles for the chromatin modifiers, Prdm3 and Prdm16, in cranial neural crest development and formation of the craniofacial skeleton. We show that these two seemingly functional redundant paralogs can act antagonistically independent of each other upstream of canonical Wnt/β-catenin signaling during chondrocyte differentiation to ensure proper spatial and temporal development of the vertebrate craniofacial skeleton.

## METHODS

### RESOURCE AVAILABILITY

#### Contact for Reagent and Resource Sharing

Futher information and requests for resources and reagents should be directed to and will be fulfilled by the Lead Contact, Kristin Artinger.

#### Materials Availability

This study did not generate new unique reagents.

#### Data and Code Availability

The RNAseq and ATACseq datasets generated in this paper will be available at NCBI Gene Expression Omnibus.

## EXPERIMENTAL MODEL AND SUBJECT DETAILS

### Zebrafish

Zebrafish were maintained as previously described [94]. Embryos were raised in defined Embryo Medium at 28.5°C and staged developmentally following published standards as described [95]. The wildtype (WT) strain used include the AB line (ZIRC). The transgenic lines used include Tg(*sox10:EGFP*) [96], Tg(*fli1:EGFP*) [78], and Tg(*7xTCF-Xla*.*Sia:NLS-mCherry*)^ia5Tg^ [77]. These transgenic lines were crossed to the various mutant backgrounds. Zebrafish mutant lines for *prdm3* and *prdm16* were generated via CRISPR-based mutagenesis as previously described [72]. The *prdm3* and *prdm16* mutant alleles used in this study (*prdm3*^*CO1005*^ and *prdm16*^*CO1006*^) are predicted frameshift mutations that interrupt the coding sequence upstream of the PR/SET domain responsible for functional histone methyltransferase activity [72]. For double mutants, *prdm3+/-* fish were bred to *prdm16+/-* fish to generate *prdm3+/-;prdm16+/-* double heterozygous animals which were intercrossed to generate *prdm3-/-;prdm16-/-* double mutants as well as all the other resulting various allelic combinatorial animals. The Institutional Animal Care and Use Committee of the University of Colorado Anschutz Medical Campus approved all animal experiments performed in this study and conform to NIH regulatory standards of care.

### Mice

*Mecom*^*tm1mik*^ (referred to as *Prdm3*^*fl/fl*^) [70], *B6(SJL)-Prdm16*^*tm1*.*1Snok*^*/J* (referred to as *Prdm16*^*fl/fl*^) (Jackson Laboratories), and *H2afv*^*Tg(Wnt-Cre)11Rth*^ (referred to as *Wnt1-Cre*^*+/Tg*^) [73] were all maintained on the C57/Bl6 background. For timed matings, *Prdm3*^*fl/fl*^ or *Prdm16*^*fl/fl*^ females were bred to *Prdm3*^*fl/+*^*;Wnt1-Cre*^*+/Tg*^ or *Prdm16*^*fl/+*^*;Wnt1-Cre*^*+/Tg*^ males, respectively. The morning a vaginal plug was detected was considered embryonic day 0.5. Embryos of matching somite numbers were used for experiments. Mice were bred and maintained in accordance with the recommendations in the Guide for the Care and Use of Laboratory Animals of the National Institutes of Health. The protocol was approved by the University of Colorado Anschutz Medical Campus’s Institutional Animal Care and Use Committee.

## METHOD DETAILS

### Genotyping

Fin clips, single whole embryos or single embryo or larva tails were lysed in Lysis Buffer (10 mM Tris-HCl (pH 8.0), 50 mM KCl, 0.3% Tween-20, 0.3% NP-40, 1 mM EDTA) for 10 minutes at 95° C, incubated with 50 ug of Proteinase K at 55°C for 2 hours, followed by inactivation of Proteinase K at 95°C for 10 minutes. Genotyping for *prdm3* and *prdm16* mutant alleles was performed as previously described [72]. For mice, tail clips from weanlings and tail clips or yolk sacs from embryos were lysed in DNA Lysis Buffer (10 mM Tris-HCl (pH 8.0), 100 mM NaCl, 10 mM EDTA (pH 8.0), 0.5% SDS) and 100 µg of Proteinase K overnight at 55°C. Genomic DNA was isolated following phenol/chloroform extraction. DNA pellets were air dried and resuspended in nuclease-free water. Genotyping for *Prdm3* and *Prdm16* alleles was performed as previously described [72]. For Wnt1-Cre PCR genotyping, the following primer sets were used: (Transgene F) 5’-CAGCGCCGCAACTATAAGAG-3’, (Transgene R) 5’-CATCGACCGGTAATGCAG-3’, (Internal Positive Control Forward) 5’-CAAATGTTGCTTGTCTGGTG-3’, and (Internal Positive Control Reverse) 5’-GTCAGTCGAGTGCACAGTTT-3’, where the transgene was detected at ∼300 bp and the internal positive control was detected at ∼200 bp.

### Inhibitor Treatments

For inhibitor treatment experiments, embryos were dechorionated at 24 hours post fertilization (hpf). The clutches were divided evenly into 2 groups, vehicle control or inhibitor treatment groups. *Prdm3* heterozygous intercrossed embryos were treated with either the Wnt inhibitor, IWR-1 (Sigma), at a final concentration of 0.75 µM or DMSO in E2 embryo water. *Prdm16* heterozygous intercrossed embryos were treated with either the Wnt activator, Gsk Inhibitor XV (CalbioChem), at a final concentration of 0.05 µM or DMSO for vehicle control in E2 embryo water. Inhibitor or vehicle treated E2 embryo water was removed after a 24 hour window (at 48 hpf developmental time) and replaced with fresh E2 embryo water. At 6 dpf, larvae were collected and fixed in 2% paraformaldehyde (PFA) and subjected to Alcian Blue/Alizarin Red staining for cartilage and bone assessment.

### Skeletal Staining

For zebrafish, Alcian Blue (cartilage) and Alizarin Red (bone) staining was performed at room temperature as previously described [97]. Briefly, 6 days post fertilization larvae were collected and fixed for one hour in 2% PFA. Following a 10-minute rinse in 100 mM Tris (pH 7.5) and 10 mM MgCl_2_, larvae were incubated in Alcian Blue solution (0.04% Alcian Blue, 80% ethanol, 100 mM Tris (pH 7.5), 10 mM MgCl_2_) overnight at room temperature. Larvae were then rehydrated and destained through a gradient of ethanol solutions 80%, 50%, 25% ethanol containing 100 mM Tris (pH 7.5) and 10 mM MgCl_2_, then bleached for 10 minutes in 3% H_2_O_2_ with 0.55% KOH at room temperature, washed twice in 25% glycerol with 0.1% KOH, then stained in Alizarin Red (0.01% Alizarin Red dissolved in 25% glycerol and 100 mM Tris (pH 7.5)) for 30 to 45 minutes at room temperature. Samples were destained in 50% glycerol with 0.1% KOH. Whole-mount and dissected and flat mounted specimens were mounted in 50% glycerol and imaged with LAS v4.4 software on a Leica stereomicroscope. High magnification images of chondrocytes were imaged with LAS v4.4 software on an Olympus compound microscope.

For mice, Alcian blue and Alizarin red staining was performed as previously described for E18.5 embryos [72, 98]. Briefly, mouse embryos were harvested at E18.5 in 1XPBS. Skin and internal organs were removed, and specimens were fixed overnight in 95% ethanol at room temperature followed by an incubation in 100% acetone for 2 days at room temperature. Embryos were then stained in Alcian Blue/Alizarin Red staining solution ((0.015% Alcian Blue, 0.05% Alizarin Red, 5% Glacial Acetic Acid and 70% ethanol) for 3 days at 37°C. Stained embryos were rinsed in water before undergoing an initial clearing in 1% KOH overnight at room temperature, followed by a gradient series of decreasing KOH concentrations and increasing glycerol concentrations. Skeletal preparations were stored and imaged in 80% glycerol on Leica stereomicroscope with the LASX v4.4 software.

### Histology

Embryos were collected at E14.5 in 1xPBS. The mandible was dissected and removed from the heads of the animals and fixed in 4% paraformaldehyde, cryoprotected in 30% sucrose solution, embedded in OCT embedding medium, sectioned to a thickness of 8 µm and mounted onto glass slides with a Leica cryostat. For staining, sections were brought to room temperature and rehydrated in 1xPBS before staining with Weigert’s iron hematoxylin, 0.05% Fast Green, and 0.1% Safranin O and mounted with Permount (Electron Microscopy Sciences). Stained sections were imaged on an Olympus compound microscope with the LASX v4.4 software.

### Fluorescence-Activated Cell Sorting (FACS) of Neural Crest Cells

At 48 hpf, *sox10:EGFP* positive embryos were stage matched and selected under a fluorescent dissecting microscope. *prdm3-/-;Tg(sox10:EGFP)* and *prdm16-/-;Tg(sox10:EGFP)* single mutant embryos were identified based on phenotype and subsequent genotyping. Two independent replicates of the *prdm3* and *prdm16* crosses were assayed. To prepare single-cell suspensions for FACS, 30-40 embryos of each genotype were dechorionated and rinsed in 1x DPBS (calcium and Mg+ free). Embryos were then dissociated in Accumax (Innovative Cell Technologies, AM-105) containing DNaseI. The samples were incubated at 31°C and agitated by pipetting every 10-15 minutes for an hour to promote cell dissociation. Following the digest, a wash solution (1x D-PBS and DNaseI) was added to stop the reaction. Cells were filtered through a 70 uM nylon mesh strainer (Fischer), centrifuged at 2000 rpm for 5 minutes at 4°C and resuspended in FACS basic sorting buffer (1 mM EDTA, 25 mM HEPES (pH 7.0), 1% FBS in 1x D-PBS). Cell suspensions were stained with DAPI (1:1000) and kept on ice. GFP-positive cells were sorted on a MoFlo XDP100 cell sorter (Beckman-Coulter) and collected in 1x DPBS. Following FACS, GFP-positive cells were then processed for RNA-sequencing or Assay for Transposase-Accessible Chromatin paired with sequencing.

### RNA-sequencing

Following FACS, GFP-positive NCCs were centrifuged briefly at 2000 rpm for 5 minutes and resuspended in TRIzol LS lysis reagent. Total RNA was extracted from sorted cells using chloroform extraction and the Direct-zol RNA miniprep kit (Zymogen) according to manufacturer’s instructions. Purified RNA quality and quantity was assessed on a High Sensitivity RNA Screen Tape (Agilent) and Infinite M200pro plate reader (Tecan). cDNA libraries were generated using the Clonetech Pico Library Prep Kit. Following library generation, sequencing was performed on an Illumina NovaSEQ 6000 system. Library construction and sequencing was performed at the University of Colorado Anschutz Medical Campus Genomics and Microarray Core Facility.

### RNA Isolation and qRT-PCR

For zebrafish, whole heads were dissected and removed from wildtype, *prdm3-/-* and *prdm16-/-* embryos at 48 hpf. 5 to 7 embryos of the same genotype were pooled and lysed in TRIzol LS (Invitrogen). Total RNA was extracted following a chloroform extraction. For mice, mandibular processes (MdP) were dissected on ice from three independent replicates of E11.5 *Prdm3*^*fl/fl*^*;Wnt1-Cre*^*+/Tg*^, *Prdm16*^*fl/fl*^*;Wnt1-Cre*^*+/Tg*^ and control (*Prdm3*^*fl/+*^*;Wnt1-Cre*^*+/Tg*^ or *Prdm16*^*fl/+*^*;Wnt1-Cre*^*+/Tg*^) embryos. The overlying ectoderm was removed by digestion in 0.25% trypsin for 10-15 mins on ice. MdPs were rinsed in 10% fetal bovine serum (FBS) for 1 minute before a quick rinse in 1xPBS. MdPs were lysed in TRIzol LS (Invitrogen) and Total RNA was isolated from these samples using the Direct-zol RNA miniprep kit (Zymogen) according to manufacturer’s instructions.

For qRT-PCR, total RNA was isolated as described above for zebrafish and mouse MdPs and (0.5-1.0 µg) was reverse transcribed to complementary DNA (cDNA) with SuperScript III First-Strand Synthesis cDNA kit (Invitrogen) for real-time semiquantitative PCR (qRT-PCR) with primers (**Supplemental Table 1**) and SYBR Green Master Mix (BioRad). Transcript levels were normalized to the reference gene, *gapdh or Actb*. Transcript abundance and relative gene expression were quantified using the 2^-ΔΔCt^ method relative to control.

### Assay for Transposase-Accessible Chromatin Paired with Sequencing

Assay for Transposase-Accessible Chromatin (ATAC)-seq was performed as previously described [99, 100]. Briefly, following tissue dissociation FAC-sorting (as described above), 25,000 GFP-positive cells per genotype were pelleted at 500g for 5 minutes at 4° C. The cell pellets were gently resuspended in cold lysis buffer (10 mM Tris-HCl, pH 7.5, 10 mM NaCl, 3 mM MgCl_2_, 0.1% v/v NP-40). Lysed cells were immediately centrifuged at 500 g for 20 minutes at 4°C. Cell nuclei pellets were resuspended in tagmentation mix (Nextera Tagmentation Buffer (Illumina), Nextera Tagmentation DNA Enzyme (Illumina). Tagmentation reaction was adjusted to accommodate 25,000 cells and incubated at 37°C for 30 minutes, mixing at 10 minute intervals. Tagmented DNA was purified using the Zymogen Clean and Concentrator kit. Libraries were amplified and indexed using NEBNext High Fidelity 2x PCR master mix. Following 11 cycles of amplification, libraries were purified with AmpureXP beads, quantified with Qubit and subjected to sequencing on the Illumina NovaSEQ 6000 system at the University of Colorado Anschutz Medical Campus Genomics and Microarray Core Facility.

### CUT&RUN paired with qRT-PCR

Cleavage Under Targets and Release Using Nuclease (CUT&RUN) was performed on wildtype whole embryos as described [101]. Briefly, ∼200 48 hpf wildtype zebrafish embryos were pooled and dissociated to a single cell suspension using Accumax and DNaseI for 1 hour with gentle pipetting to agitate the tissue every 10 minutes. Following dissociation, a wash solution containing 1xPBS and DNaseI was added to stop the reaction. Cells were passed through a 70 µm filter and counted. 500,000 cells were incubated on activated Concanavalin A conjugated paramagnetic beads (Epicypher) for 10 minutes at room temperature. Cell bound beads were resuspended in antibody buffer (20 mM HEPES, pH 7.5; 150 mM NaCl; 0.5 mM Spermidine (Invitrogen); 1x Complete-Mini Protease Inhibitor tablet (Roche); 0.01% Digitonin (Invitrogen); 2mM EDTA) and incubated in the corresponding antibodies: IgG, Prdm3/Evi1, Prdm16, and H3K27ac overnight with rotation at 4°C. Next day, cells were washed in Digitonin Buffer wash solution (20 mM HEPES, pH 7.5; 150 mM NaCl; 0.5 mM Spermidine (Invitrogen); 1x Complete-Mini Protease Inhibitor tablet (Roche); 0.01% Digitonin (Invitrogen)) twice and then incubated with pAG-MNase (Epicypher) for 10 minutes at room temperature. Following another 2 washes in Digitonin Buffer, 1 µl of 100 mM CaCl_2_ was added to each sample and incubated at 4°C for 2 hours. This digestion reaction was stopped with Stop Buffer (340 mM NaCl, 20 mM EDTA, 4 mM EGTA, 50 µg/ml RNaseA, and 50 µg/ml Glycogen) and another incubation for 10 minutes at 37°C. DNA fragments were purified using a DNA Clean and Concentrator Kit (Zymogen). For qRT-PCR, eluted CUT&RUN fragmented DNA was amplified using the NEBNext Ultra II DNA Library Prep Kit as per manufacturer’s instructions. qRT-PCR was performed with primers designed to flank putative Prdm3 and Prdm16 binding motifs at promoter regions of Wnt/β-catenin targets genes or negative control genes with no binding sites (*gapdh*) and SYBR Green Master Mix (BioRad). The fold enrichment for the abundance of Prdm3, Prdm16, or H3K27ac of those amplified regions was calculated and normalized relative to the IgG control and averaged across 3 different experiments.

### Whole-Mount Immunofluorescence

Zebrafish larvae were collected at the indicated timepoints and fixed in 4% paraformaldehyde overnight at 4°C. Following fixation, embryos were washed in 1x Phosphate Buffered Saline (PBS, pH 7.3) with 1% TritonX-100 three times for 10 min at room temperature. For antigen retrieval, embryos were incubated in 1 ug/ml Proteinase K diluted in 1x PBS with 1% Triton X-100 for 20 minutes at room temperature. Following proteinase K treatment, embryos were incubated in 4% PFA for 15 minutes at room temperature, then washed 3 more times in 1x PBS with 1% Triton X-100 for 10 minutes at room temperature. Embryos were blocked at room temperature for 1 hour in blocking solution containing 10% normal goat serum and 1% bovine serum albumin in 1x PBS. Samples were incubated in primary antibodies diluted in blocking solution [anti-acetylated α-tubulin, anti-phosphorylated Y489 β-catenin, Rhodamine phalloidin, Wheat Germ Agglutinin], overnight at 4°C. Following primary antibody incubation, samples were thoroughly washed in PBS with 1% Triton X-100 before incubating with corresponding secondary antibodies overnight at 4°C, then washed again thoroughly in 1x PBS with 1% Triton X-100 before incubation with DAPI diluted in 1x PBS for 1 hour at room temperature. Samples were quickly washed in PBS with 1% Triton X-100 before mounted in Vectashield mounting media (Invitrogen) on glass slides. Embryos were imaged on a Leica TCS SP8 confocal microscope. Images were processed in LASX software and ImageJ.

## QUANTIFICATION AND STATISTICAL ANALYSIS

### RNA-sequencing Bioinformatics Analysis

Following trimming and read alignment, paired-end reads were mapped to the zebrafish genome (danRer11) assembly using TopHat [102, 103]. Differential expression between mutant and wildtype was calculated using Cufflinks [103]. Gene expression was expressed in fragments per kilobase of transcript per million mapped reads (FPKM). Complete RNA-seq datasets will be available at the Gene Expression Omnibus repository. Normalized counts were converted to z-scores for plotting heatmaps using the pheatmaps R package (https://cran.r-project.org/web/packages/pheatmap/index.html). Gene lists were analyzed for functional annotation using GO enrichment analysis based on the PANTHER Classification System [104-107].

### ATAC-seq Bioinformatics Analysis

Sequencing reads were aligned to the zebrafish genome (danRer11) using the HISAT2 mapping tool [108]. The specific CTGTCTCTTATA barcode was removed using Cutadapt before mapping. Multiple-mapped and duplicated reads were filtered out by using the modified assign_multimappers.py and picard MarkDuplicates programs, respectively. The resulting clean de-duplicated well paired reads were then used for inferring the fragment tags and generating the final BAM files. Peaks were called using the MACS2 peak calling program [109]. Annotation of peaks corresponding to TSS/promoter, intergenic, intronic and TES locations was carried out using HOMER (v4.11) annotatePeaks.pl script [110]. DeepTools (v2.0) was used to generate bigWig coverage files for visualization [111]. Transcription factor Occupancy prediction by Investigation of ATAC-seq Signal (TOBIAS) was performed following the standard pipeline and workflow [80]. Complete ATAC-seq datasets will be available in the GEO repository.

### Image Quantification

To quantify chondrocyte organization from Alcian stained and dissected flat mounts, high magnification images of cartilage elements were imported into ImageJ. An angle was drawn from the center of three adjacent chondrocytes moving along the cartilage element in the direction of chondrocyte growth. Angles were measured and averaged across at least 5 cartilage elements (namely the posterior ceratobranchial cartilages) per individual. To quantify chondrocyte cell orientation, individual chondrocytes in the developing palatoquadrate were divided into quadrants in ImageJ. The positioning of acetylated α-tubulin puncta was indicated as either 0°, 90°, 180°, or 270° in the direction of growth of the palatoquadrate, i.e. anteriorly toward the jaw joint junction with the Meckel’s cartilage. Positioning was tracked through z-stack images and the number of acetylated α-tubulin puncta in each quadrant were tabulated for each individual and normalized to the total number of cells analyzed for that individual. Total counts for at least 5 individuals per were averaged across genotypes and plotted as circular graphs. Nuclear β-catenin puncta (Phosphorylated Y489 β-catenin) were quantified and tracked across 10 chondrocytes in the palatoquadrate through z-stack images for one individual. The number of puncta was averaged for each individual and at least 5 individuals were analyzed per genotype.

For quantifying chondrocyte cell area in mouse Meckel’s cartilage, the area of 200 cells across 4-5 sections anterior to posterior through the tissue were measured and averaged across 3 individuals per genotype. For cell numbers, cells were counted in a designated region of tissue area across 4-5 sections anteriorly to posteriorly throughout the tissue and averaged across 3 individuals per genotype.

### Statistical Analysis

Data shown are the means ± SEM from the number samples or experiments indicated in the figure legends. All assays were repeated at least three times with independent samples. *P* values were determined with Student’s *t* tests.

## Acknowledgements

We thank members of the Artinger laboratories for their thoughtful discussions on this project and Dr. Richard Dorsky for the *Tg(7xTCF-Xla*.*Siam:NLS-mCherry)* line. We thank Jeremy Dasen, David Schoppik, and Kristen D’Elia at the New York University for their feedback and insightful comments on this project. Special thanks to David Clouthier for mouse craniofacial anatomy consultation. We thank the Denver zebrafish community, and the zebrafish and mouse facility staff for excellent animal care. Funding was provided by the National Institute of Dental and Craniofacial Research (R01 DE024034 to K.B.A. and F32 DE029099 to L.C.S.) and the National Institute of Neurological Disorders and Stroke (P30 NS048154 to the UC Anschutz Medical Campus zebrafish core facility). The bioinformatics computing resources were funded by National Cancer Institute (P30CA046934 to the University of Colorado Cancer Center).

## Author Contributions

Conceptualization, L.C.S and K.B.A.; Methodology, L.C.S. and K.B.A.; Investigation, L.C.S. and E.S.L; Formal Analysis, L.C.S., H.M.K., J.C.C., K.J.; Writing – Original Draft, L.C.S. and K.B.A.; Writing – Review & Editing, L.C.S., E.S.L., J.C.C. and K.B.A.; Funding Acquisition, L.C.S. and K.B.A.; Resources, S.G. and M.K.; Supervision, K.B.A.

